# A multi-institutional investigation of psilocybin’s effects on mouse behavior

**DOI:** 10.1101/2025.04.08.647810

**Authors:** Odilia D. Lu, Katrina White, Kendall Raymond, Christine Liu, Alexandra S. Klein, Nathaniel Green, Sam Vaillancourt, Austin Gallagher, Lena Shindy, Alyssa Li, Kira Wallquist, Ruoxian Li, Mila Zou, Austen B. Casey, Lindsay P. Cameron, Matthew B. Pomrenze, Vikaas Sohal, Mazen A. Kheirbek, Andrea M. Gomez, Stephan Lammel, Boris D. Heifets, Robert Malenka

**Author notes:** These authors contributed equally (Odilia D. Lu, Katrina White, Kendall Raymond, Christine Liu, Alexandra S. Klein, Nathaniel Green). Correspondence to: Robert Malenka, M.D., Ph.D., Department of Psychiatry and Behavioral Sciences, Lorry Lokey Stem Cell Research Building, 265 Campus Drive, Room G1021, Stanford, California 94305-5453.

## Abstract

Studies reporting novel therapeutic effects of psychedelic drugs are rapidly emerging. However, the reproducibility and reliability of these findings could remain uncertain for years. Here, we implemented a multi-institutional collaborative approach to define the robust and replicable effects of the psychedelic drug psilocybin on mouse behavior. Five laboratories performed the same experiments to test the acute and persistent effects of psilocybin (2 mg/kg, IP) on various behaviors that psychedelics have been proposed to affect, including anxiety-related approach-avoidance, exploration, sociability, depression-related behaviors, fear extinction, and social reward learning. Through this coordinated approach, we found that psilocybin had several robust and replicable acute effects on mouse behavior, including increased anxiety- and avoidance-related behaviors and decreased fear expression. Surprisingly, however, we found that psilocybin did not have replicable effects 24 hours post psilocybin administration on reducing anxiety- and depression-like behaviors or facilitating fear extinction learning. Additionally, we were unable to observe psilocybin-induced alterations in social preference or social reward learning. Overall, our comprehensive characterization of psilocybin’s acute and persistent behavioral effects using ∼200 total male and female mice per experiment spread across five independent labs demonstrates with unique certainty several acute drug effects and suggests that psilocybin’s persistent effects in mice may be more modest and inconsistent than previously suggested. We believe this unusual multi-laboratory, highly coordinated research effort serves as a model for facilitating the generation of replicable results and consequently will reduce efforts based on unreliable and spurious results.

## Introduction

After decades of limited research, there is now renewed interest in studying psychedelic drugs. A single or few doses of various psychedelic drugs including psilocybin^1–8^, MDMA^9,10^, LSD^11,12^, and ibogaine^13–15^, appear to elicit rapid and enduring therapeutic effects for a wide range of neuropsychiatric disorders, including anxiety^4,5,11–13,15^, depression^1–5,8,13,15^, substance use disorder^6,7,14^, and post-traumatic stress disorder^9,10,15^. These preliminary clinical findings have generated great excitement in both the lay and scientific press because they suggest that psychedelic drugs may have unique therapeutic properties that could dramatically transform psychiatry. Indeed, three psychedelic drugs (MDMA, LSD, and psilocybin) have been granted Breakthrough Therapy status by the Food and Drug Administration^16–19^.

Advancing our understanding of the uses for, and mechanisms of, psychedelic drugs requires that clinical and basic science findings be robust (i.e., the effect size is large) and replicable (i.e., the same results can be obtained by different groups performing the same study). However, the field currently struggles to demonstrate both robustness and replicability. For example, psilocybin, which has arguably received the greatest attention because of its promise for treating depression, failed to produce stronger antidepressant effects than the selective serotonin reuptake inhibitor escitalopram^20^. Some have questioned whether its effects surpass those expected from placebo^21^. Furthermore, inconsistent findings are evident, both in terms of psilocybin’s behavioral outcomes^22–26^ and therapeutic mechanisms of action in mice^23,27,28^. Given the potential for psychedelic drugs like psilocybin to reshape psychiatric medicine, efficiently advancing our understanding of the robust and replicable effects of psychedelics on behavior is essential.

To facilitate the efficient collection of replicable results in psychedelic science, we believe that a new research approach is necessary. Current norms in academic research, in which scientists independently investigate identical or similar questions, can drive differences in results likely due in part to slight differences in methodology^29,30^. For example, in the context of psychedelic drug research, several studies have sought to determine whether psilocybin has presumptive “antidepressant” effects in rodents. While some studies demonstrate significant effects in the forced swim test^24,25,31^, other studies report null results^22,23,26^. Without highly consistent methodology and extensive communication between individual investigators, we anticipate that discrepancies in psilocybin’s effects across independent studies will become increasingly difficult to resolve. Ultimately, the lack of collaboration and coordination among laboratories pursuing similar objectives will hinder our understanding of psilocybin’s replicable effects and thereby slow progress towards understanding its potential therapeutic mechanisms of action.

In this study, we used a novel, collaborative approach to define the robust and replicable behavioral effects of the psychedelic drug psilocybin in mice. Specifically, we established a consortium of five labs (termed the Psychedelic Bay Area Animal Neuroscience Consortium; Psy-BAANC), in which each lab performed the same psilocybin behavioral experiments, aligning on dose (2 mg/kg, IP), strain (C57BL/6J mice), age (∼10-14 weeks old), sexes used (both male and female mice), experimental protocols, and timepoints of testing. We characterized both acute (∼10 minutes after drug administration) and persistent (24 hours after drug administration) effects of the drug and focused on testing behaviors that classic psychedelics, including psilocybin, have been reported to affect.

## Results

### Psilocybin robustly increases head twitch responses

To identify the replicable effects of psilocybin on behavior, it was critical to select an appropriate dose. We chose 2 mg/kg because this dose: (1) triggers head twitch responses in mice which are commonly considered a correlate of hallucinogenic activity^32,33^, (2) is low enough to avoid gross motor changes^34^, and (3) is similar to doses used in other preclinical studies^22,23,27,35–39^. After confirming the concentration and stability of our drug preparation (**Extended Data Fig. 1**), each lab tested drug efficacy by monitoring head twitch responses for 20 minutes after administering psilocybin intraperitoneally (**Fig. 1a**). Psilocybin consistently increased head twitch responses, both in each individual lab (**Fig. 1b**) and when summarizing data across all labs (**Fig. 1c**). Modest sex differences were evident, with female mice generally exhibiting more psilocybin-induced head twitches than male mice. Consistent with previous reports^22,36^, a detailed time course analysis suggested that psilocybin-induced head twitches were most prominent in the first ten minutes after drug administration (**Fig. 1d, 1e**).

**Figure 1.**
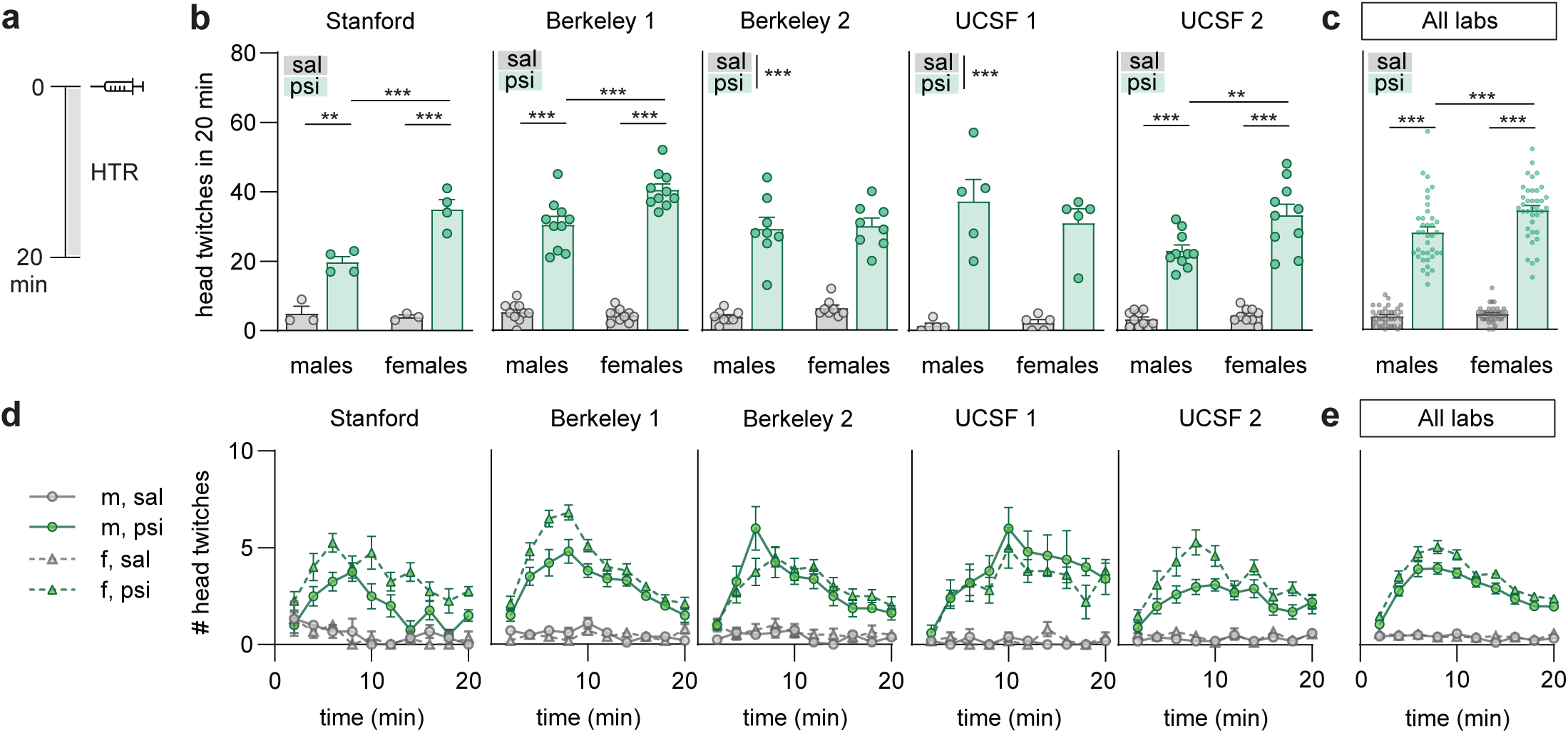
Psilocybin increases head twitch responses in mice across labs. **(a)** Experimental timeline to assess acute effects of psilocybin (2 mg/kg, IP) on head twitch response (HTR). **(b)** Number of head twitches observed in the first 20 minutes after psilocybin administration for male and female mice. Each graph shows data from a different lab. (sal: saline treatment; psi: psilocybin treatment) **(c)** Data in panel b pooled together. **(d)** Time course analysis of head twitches (2-min time bins) for male saline-treated mice (m, sal), male psilocybin-treated mice (m, psi), female saline-treated mice (f, sal), and female psilocybin treated mice (f, psi). **(e)** Data in panel d pooled together. Data shown as mean+SEM; **p<0.01, ***p<0.001. Statistics and significance visualization are explained in **Methods**. Statistical reports for all analyses are provided in **Supplementary Table 2**.

### Psilocybin acutely increases anxiety-like behaviors with no persistent anxiolytic effects

Having confirmed the behavioral activity of psilocybin within the setting of each individual lab, we assessed the acute and persistent effects of the drug on behaviors that psychedelics are thought to alter, including behaviors related to anxiety^22,35,40^, exploration^41,42^, sociability^43,44^, depression^23,24,27,28,31^, fear extinction learning^37,39,45,46^, and social reward learning^47,48^. To assess psilocybin’s acute effects on anxiety-related approach-avoidance behaviors, we first performed the open field test^49^ (**Fig. 2a**). Four of five labs found psilocybin significantly reduced time in the center, with the Stanford lab observing a strong trend in the same direction (**Fig. 2b**). Pooling data from all labs revealed a strong effect of the drug despite the great variability in individual measurements (**Fig. 2c**). We next tested the acute effects of psilocybin in the elevated plus maze; a second anxiety-related behavioral test^50^ (**Fig. 2d**). Three of five labs found that psilocybin-treated mice spent significantly less time in the open arms, with the other two labs generally observing effects in the same direction (**Fig. 2e**). Pooling results from all labs again revealed a highly significant anxiogenic effect of psilocybin despite great variability in individual measurements (**Fig. 2f**).

**Figure 2.**
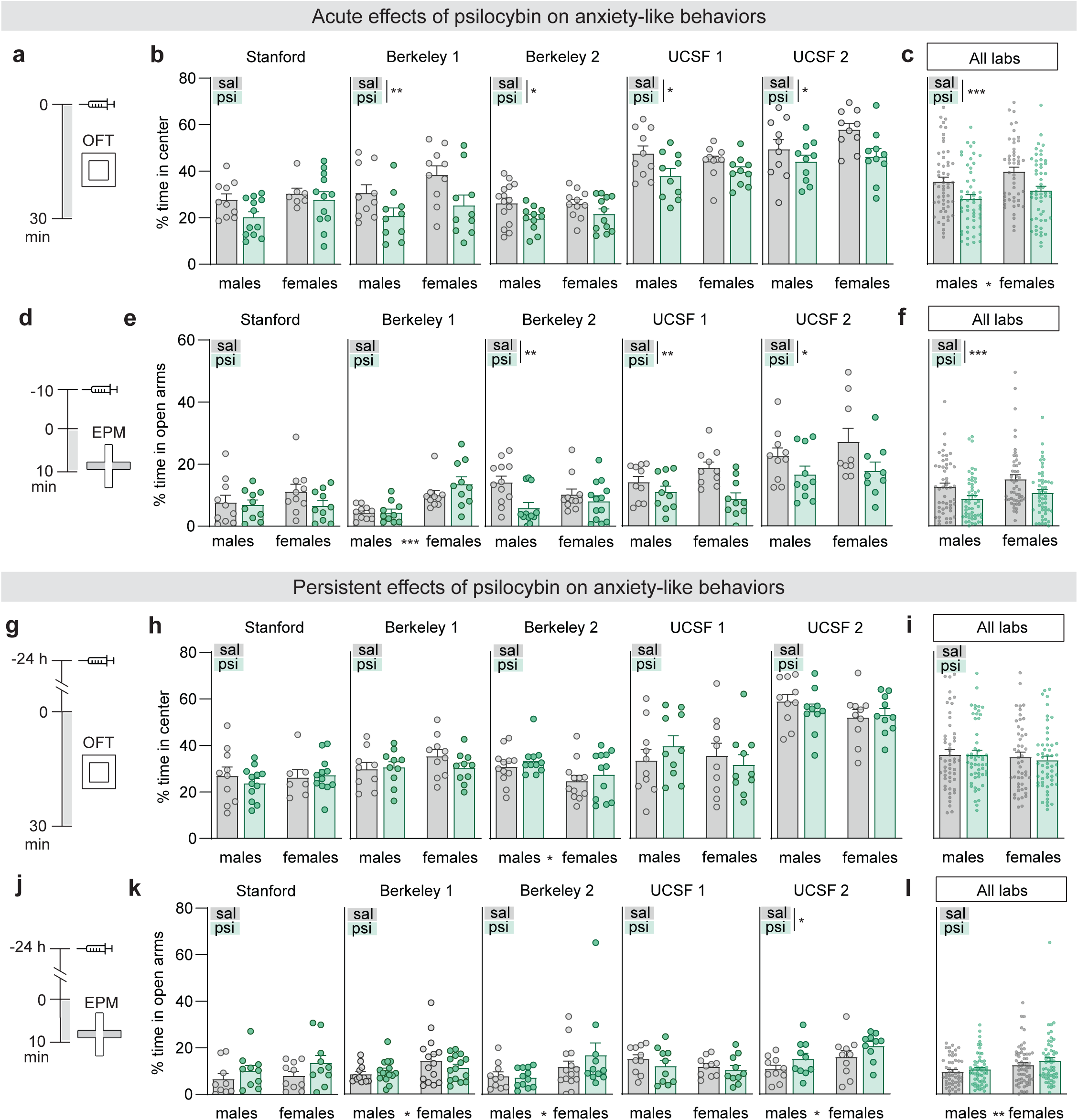
Psilocybin acutely increases anxiety-like behaviors but has no replicable persistent effects. **(a)** Experimental timeline to test acute effects of psilocybin on anxiety-like behaviors in the open field test (OFT). **(b)** Percentage of time mice spent in the center of the open field for each individual lab. **(c)** Data in panel b pooled together. **(d)** Experimental timeline to test acute effects of psilocybin on anxiety-like behaviors in the elevated plus maze (EPM). **(e)** Percentage of time mice spent in the open arms of the elevated plus maze for each individual lab. **(f)** Data in panel e pooled together. **(g-l)** As in panels a-f, but for experiments testing the long-lasting effects of psilocybin on anxiety-like behaviors (24 hours after drug administration). Data shown as *p<0.05, **p<0.01, ***p<0.001. Statistics and significance visualization are explained in **Methods**. Statistical reports for all analyses are provided in **Supplementary Table 2**.

Having found that psilocybin acutely increases anxiety-related avoidance behaviors in a replicable manner across labs, we performed the same assays 24 hours after psilocybin administration. In the open field test (**Fig. 2g**), no individual labs detected a persistent effect of psilocybin on center time (**Fig. 2h**), and no effect was revealed when results from all labs were pooled (**Fig. 2i**). Similarly, in the elevated plus maze (**Fig. 2j**), four of five labs found no persistent effects of psilocybin on the time spent in the open arms (**Fig. 2k**), and again no effect was revealed when all results were pooled (**Fig. 2l**). We also examined overall locomotion in the open field and observed no acute or persistent effects of psilocybin (**Extended Data Fig. 2**).

### Psilocybin acutely reduces novel object exploration with no persistent effects

Avoidant coping strategies are frequently observed in anxiety^51^. Therefore, we next tested whether psilocybin influenced avoidance of novel objects (**Fig. 3a**). Although there was variability in results between labs, with three labs finding that psilocybin acutely decreased novel object exploration and the other two labs reporting inconsistent effects (**Fig. 3b**), pooling results from all labs revealed a significant decrease in females and a nonsignificant effect in the same direction in males (**Fig. 3c**).

**Figure 3.**
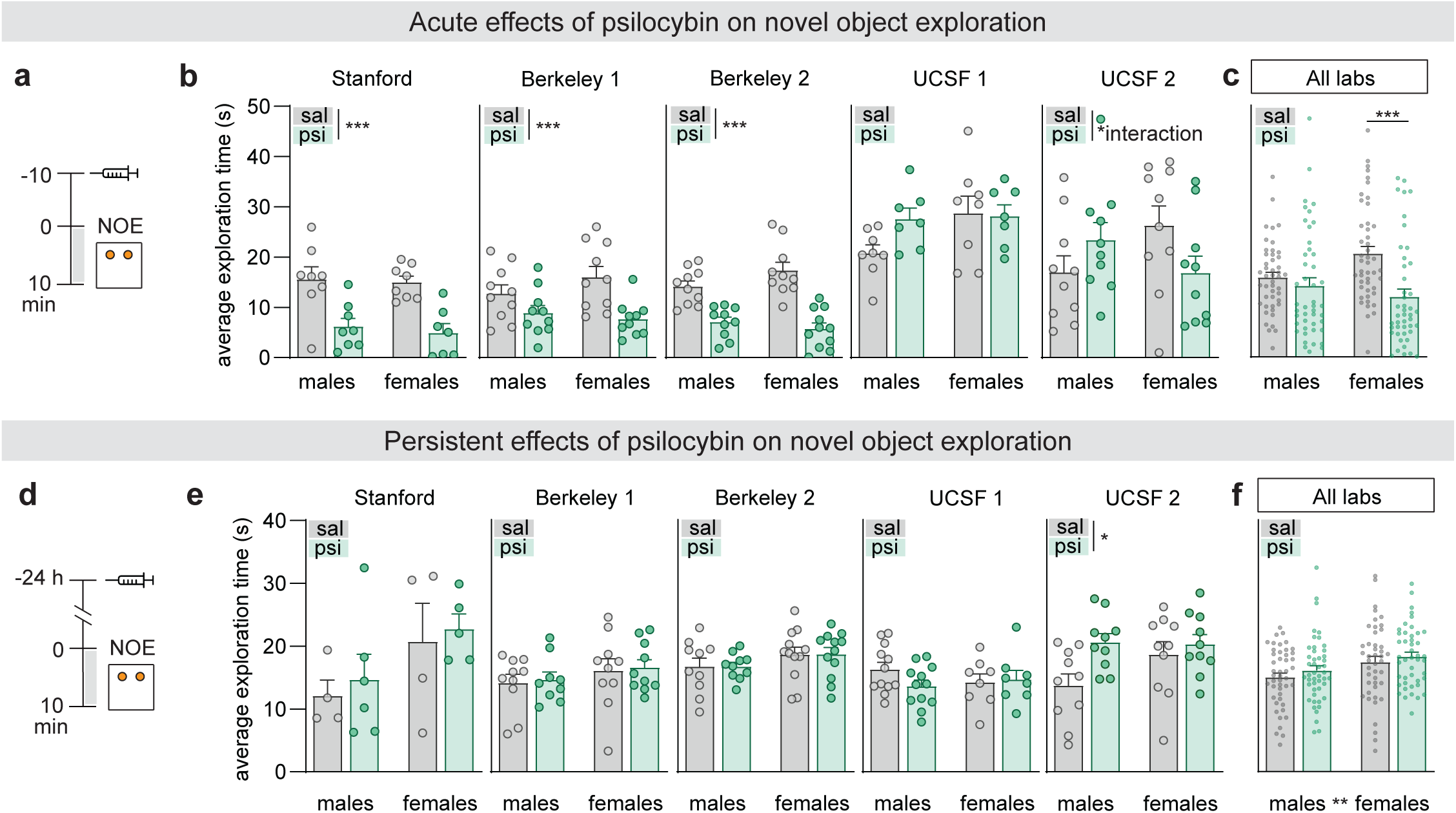
Psilocybin acutely reduces novel object exploration but has no replicable persistent effects. **(a)** Experimental timeline to test acute effects of psilocybin on novel object exploration (NOE). **(b)** Average mouse exploration time of the novel objects for each lab. **(c)** Data in panel b pooled together. **(d-f)** As in panels a-c, but for experiments testing the persistent effects of psilocybin on NOE (24 hours after drug administration). Data shown as mean+SEM; *p<0.05, ***p<0.001. Statistics and significance visualization are explained in **Methods**. Statistical reports for all analyses are provided in **Supplementary Table 2**.

We also tested the persistent effects of psilocybin on novel object exploration by monitoring behavior in this test 24 hours after drug administration (**Fig. 3d**). Four of five labs found no effects of psilocybin on object exploration at this timepoint with the remaining lab reporting an increase in exploration (**Fig. 3e**). As expected from the individual labs’ results, pooled results revealed no persistent effects of psilocybin (**Fig. 3f**).

### Psilocybin has no acute or persistent effects on social preference

Psychedelics have been reported to influence social interactions^43,44^. Therefore, we next examined psilocybin’s acute and persistent effects on social approach/avoidance using the three-chamber social interaction test^52^ (**Fig. 4a**). All labs found that subjects spent more time exploring the cup containing a conspecific, with individual lab social preference indexes ranging on average from as low as ∼0.2 to as high as ∼0.6. However, no lab observed either an acute (**Fig. 4b-d**) or persistent (**Fig. 4e-h**) effect of psilocybin on social preference.

**Figure 4.**
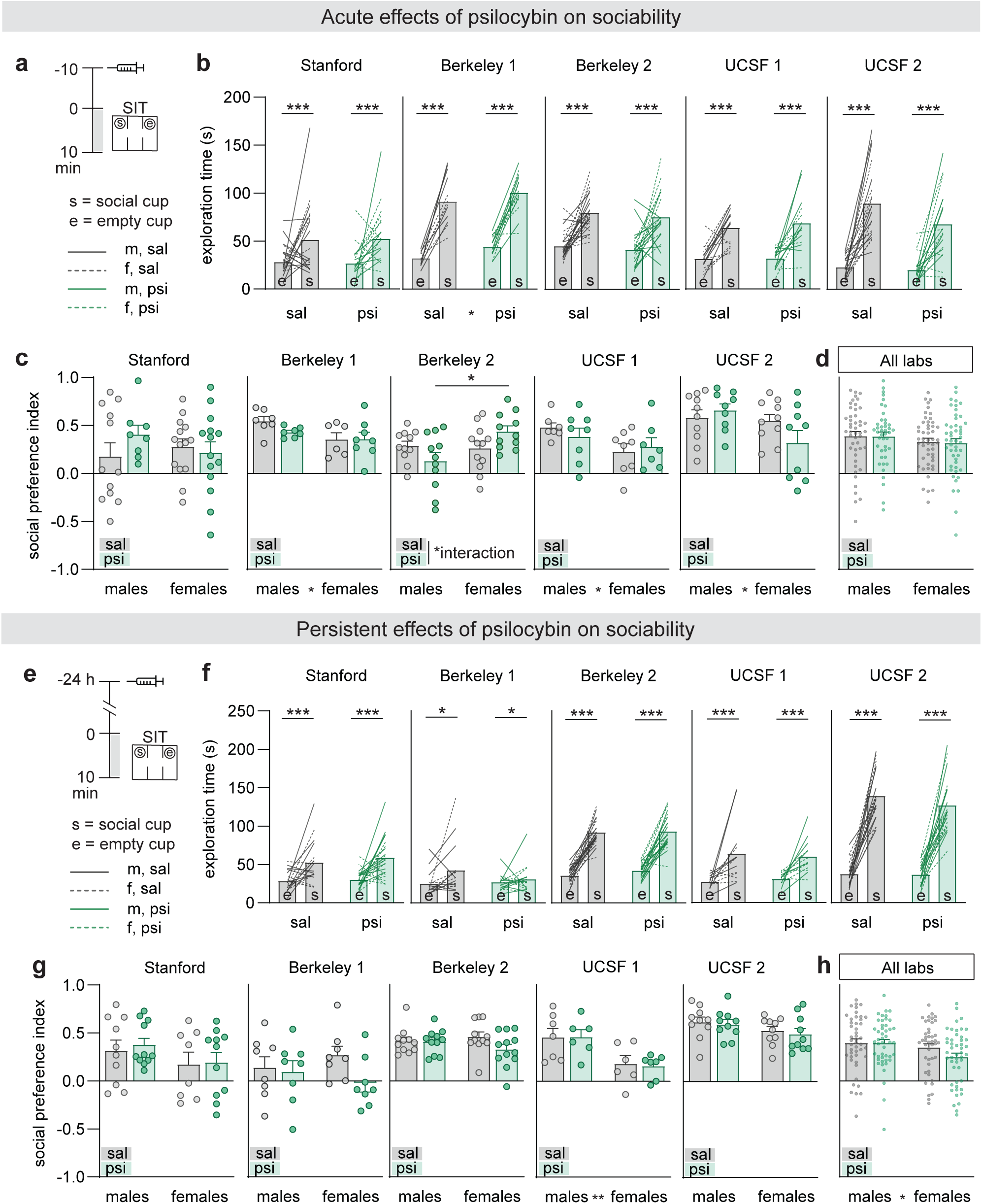
Psilocybin has no acute or persistent effects on social preference. **(a)** Experimental timeline to test the acute effects of psilocybin on social preference in the 3-chambered social interaction test (SIT). **(b)** Raw exploration time data for each lab, showing a significant main effect of cup type on exploration time (e: empty cup, s: social cup). **(c)** Social preference index calculated from data in b for each lab. **(d)** Data in panel c pooled together. **(e-h)** As in panels a-d, but for experiments testing the persistent effects of psilocybin on social preference in SIT (24 hours after drug administration). Data shown as mean+SEM; *p<0.05, **p<0.01, ***p< 0.001. Statistics and significance visualization are explained in **Methods**. Statistical reports for all analyses are provided in **Supplementary Table 2**.

### Psilocybin does not have replicable antidepressant-like effects

Psilocybin has shown promise as a potential treatment for depression^1–3,23,27,28,31^, motivating us to examine psilocybin’s effects in the commonly used forced swim test (FST)^53^. Specifically, we replicated a protocol in which the FST is performed 1 day pre- and 1 day post-drug administration (**Fig. 5a**) because several psychedelic drugs and analogs have been reported to elicit anti-depressant activity as demonstrated by reduced immobility in the post-drug assay^28,54–56^. While this repeated FST design has been reported to induce an increase in immobility on the second day (often interpreted as a “depression-like phenotype”)^28^, this effect was not replicable, as only two labs observed an increase in immobility during the second FST following saline administration (**Fig. 5b**). Furthermore, psilocybin’s antidepressant-like effects in this assay were not replicable, as four out of five labs found no significant effects of psilocybin on immobility in the second FST following psilocybin administration (**Fig. 5c**), and pooled results from all labs did not yield significant effects (**Fig. 5d**). Additional experiments replacing the FST with the tail suspension test (TST)^57^ (**Fig. 5e**) yielded a consistent increase in immobility across days in all five labs (**Fig. 5f**), but again, no antidepressant-like effects of psilocybin in this assay were observed by any lab (**Fig. 5g**) or in the pooled data (**Fig. 5h**). TST experiments performed in naïve mice also revealed no significant main effects of psilocybin (**Extended Data Fig. 3**). Taken together, these results demonstrate that psilocybin does not cause reliable changes in two commonly used assays of anti-depressant activity in mice.

**Figure 5.**
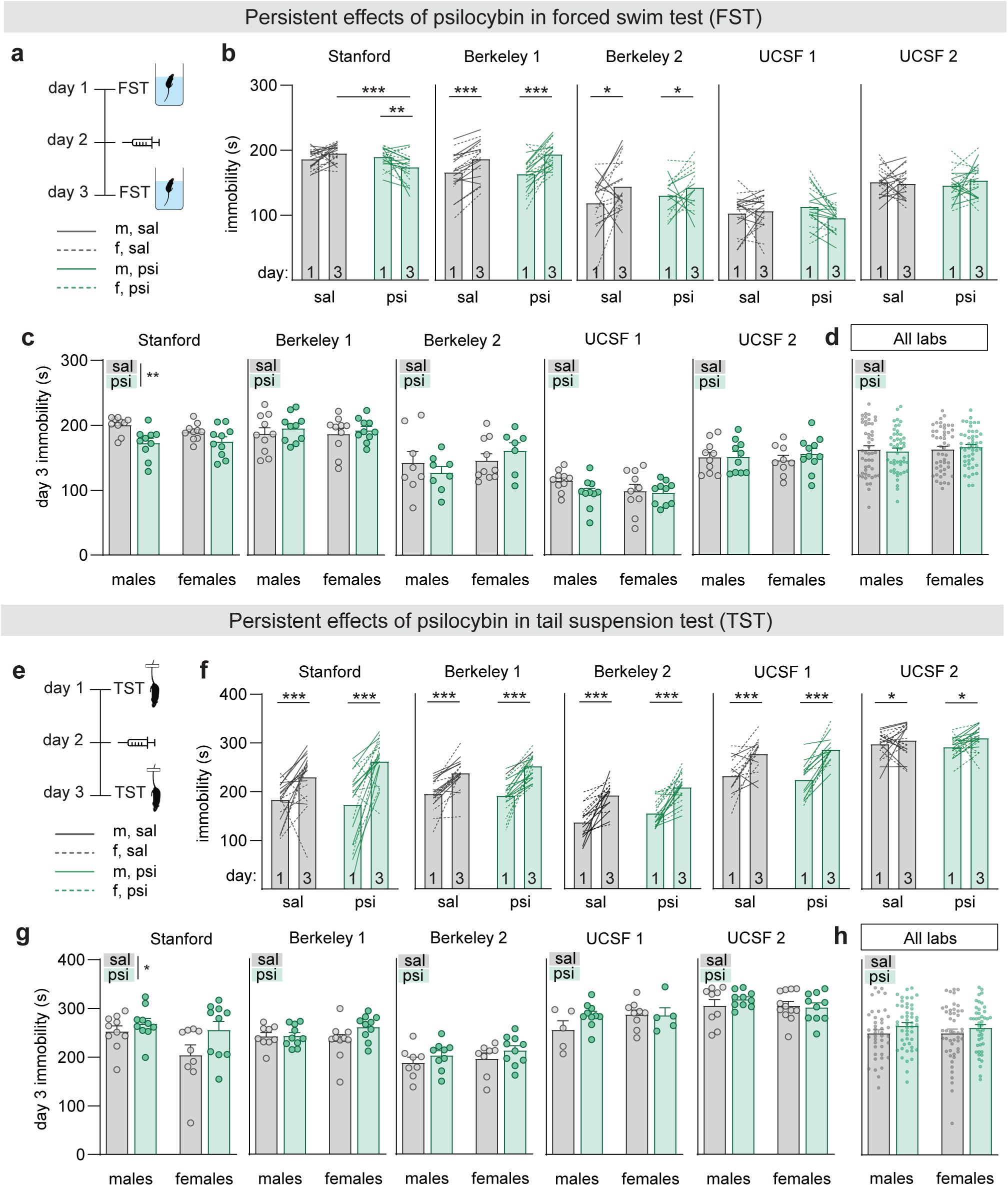
Psilocybin has no replicable persistent effects on depression-related behaviors. **(a)** Experimental procedures to test the persistent effects of psilocybin on depression-like behaviors in the repeated forced swim test (FST). **(b)** Raw immobility scores for each lab, comparing day 1 to day 3 immobility for saline (sal) versus psilocybin (psi) groups. **(c)** Day 3 immobility scores (1 day after drug administration). **(d)** Data in panel c pooled together. **(e-h)** As in panels a-d but using the tail suspension test (TST) instead of the FST. Data shown as mean+SEM; *p<0.05, **p<0.01, ***p<0.001. Statistics and significance visualization are explained in **Methods**. Statistical reports for all analyses are provided in **Supplementary Table 2**.

### Acute psilocybin reduces fear expression but does not produce replicable persistent effects on fear extinction

Psilocybin has been reported to enhance fear memory extinction potentially through its plasticity-promoting effects^37,39^. To examine the replicability of these results, we performed a standard cued fear conditioning experiment during which psilocybin was administered either immediately before or after retrieval. Administration of psilocybin before fear memory retrieval (**Fig. 6a**) caused a robust decrease in freezing during retrieval in all labs (**Fig. 6b, c**). However, no persistent effects of psilocybin on fear extinction were observed 24 and 48 hours after drug administration in four out of five labs (**Fig. 6b**). While psilocybin had a small but significant persistent effect on facilitating fear extinction in the pooled results (**Fig. 6c**), this effect was driven by a single lab (UCSF 1) finding a large effect size. Administration of psilocybin after fear memory retrieval (**Fig. 6d**) produced no persistent effects on fear extinction in four out of five labs (**Fig. 6e**) and in the pooled data (**Fig. 6f**).

**Figure 6.**
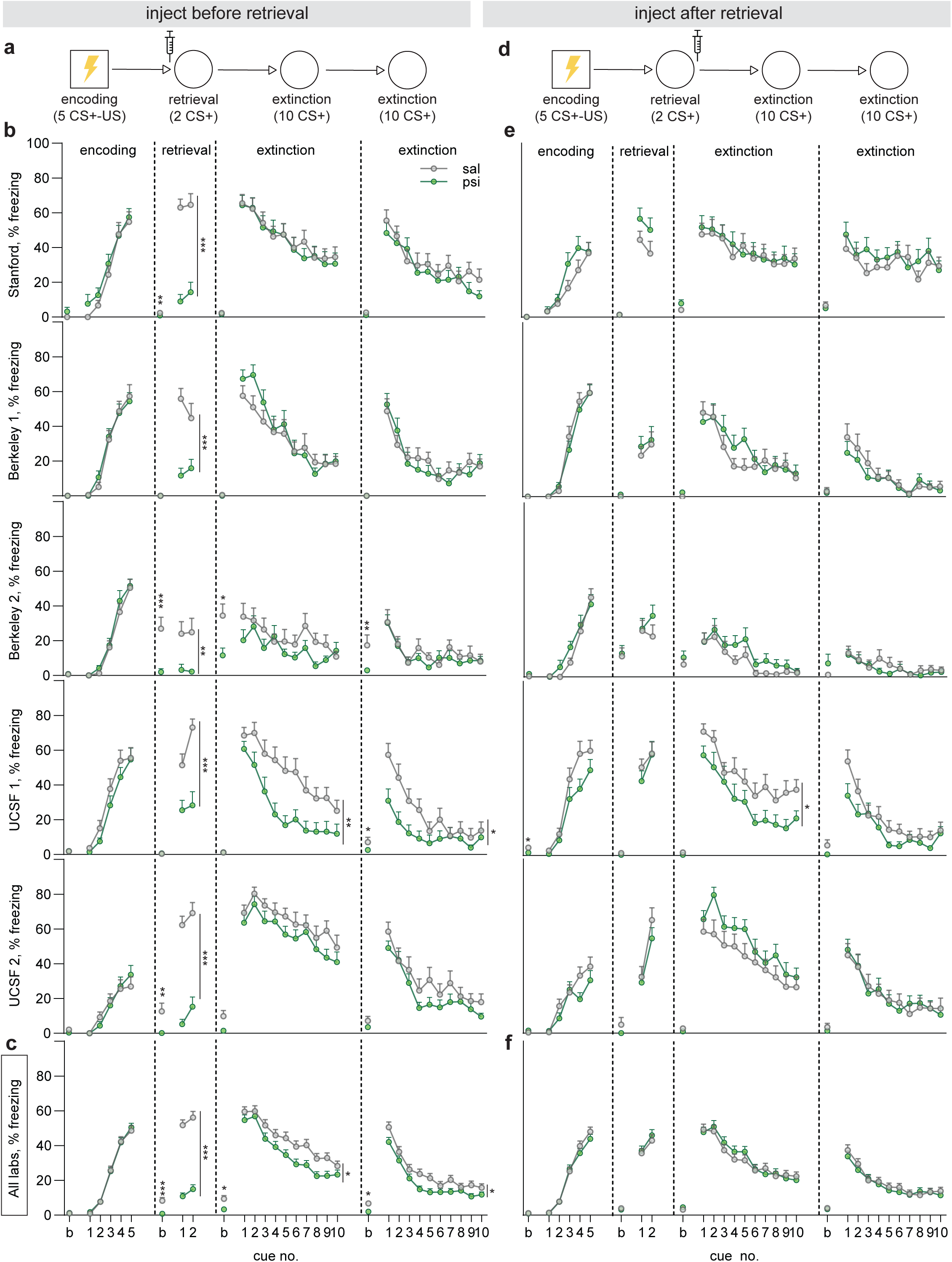
Psilocybin acutely reduces fear expression but has no replicable persistent effects on fear extinction. **(a)** Experimental timeline to test whether psilocybin injected directly before fear memory retrieval facilitates fear extinction. **(b)** Percentage of time mice spent freezing during baseline (b), and for each cue during encoding, retrieval, extinction day 1, and extinction day 2. Each graph is data from an individual lab. **(c)** Data in panel b pooled together. **(d-f)** As in a-c, but for experiments testing whether psilocybin injected after fear memory retrieval facilitates fear extinction. Data shown as mean+SEM; *p<0.05, **p<0.01, ***p<0.001. Statistics and significance visualization are explained in **Methods**. Statistical reports for all analyses are provided in **Supplementary Table 2**.

### Psilocybin does not reopen a social reward learning critical period

Recently, it has been suggested that a broad array of psychedelic drugs can reopen a social reward learning critical period^47^, which was defined by the ability of young (∼21-80 days old) but not adult (>90 days old) mice to exhibit modest preference for a context in which they previously had social interactions with cagemates for 24 hours over a context in which they were socially isolated for 24 hours^47,48^. In that study, psilocybin facilitated social reward learning in mice ∼100 days old even when administered 2 weeks prior to the conditioning procedure^47^. To test the replicability of this phenomenon, we used a simplified social reward learning protocol (**Fig. 7a**), designed to mimic a conditioned place preference assay, which is commonly used to test the rewarding properties of drugs of abuse^58,59^. Using this assay, three of five labs found that juvenile (34-41 days old) but not adult (88-106 days old) mice exhibited a place preference for the context in which they had previously experienced social interactions (**Fig. 7b-e**), thereby suggesting the presence of a critical period for social reward learning^47,48^. Although we used a slightly different assay for social reward learning, it is noteworthy that the magnitude of the social preference in the juvenile and adult mice that we observed was virtually identical to that previously reported (Juveniles = ∼1.2, Adults = ∼1.0)^47,48^. The remaining two labs did not observe robust social reward learning in juvenile mice (**Extended Data Fig. 4**).

**Figure 7.**
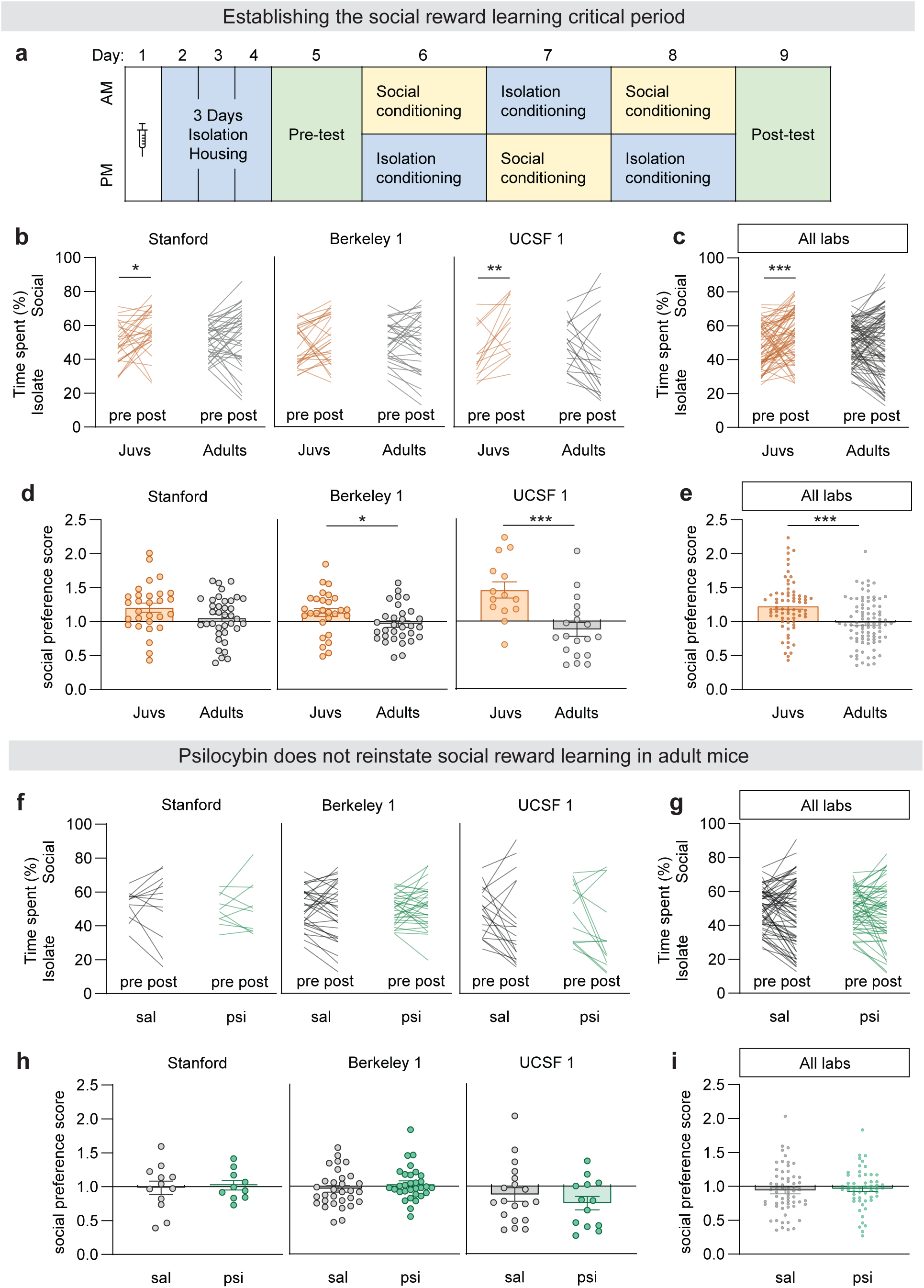
Psilocybin does not reopen social reward learning critical periods. **(a)** Social conditioned place preference test graphical methods. **(b)** Percentage of time mice spent in the social context during pre-test versus post-test, for juvenile and adult mice, in three individual labs. Percentages over 50% indicate that the mouse spent more time in the social context, and percentages under 50% indicate that the mouse spent more time in the isolation context. **(c)** Data in panel b pooled together. **(d)** Social preference score for juvenile and adult mice in each individual lab. **(e)** Data from panel d pooled together. **(f-i)** As in b-e, but comparing saline-treated adult mice (gray) versus psilocybin-treated adult mice (green). Data shown as mean±SEM; *p<0.05, **p<0.01. Statistics and significance visualization are explained in **Methods**. Statistical reports for all analyses are provided in **Supplementary Table 2**.

To test whether psilocybin reopens this social reward learning critical period, the three labs that observed social reward learning in juvenile mice administered saline or psilocybin to adult mice (88-106 days old) four days before the start of the social conditioned place preference protocol. No lab observed social reward learning following either saline or psilocybin treatment (**Fig. 7f-i**), indicating that psilocybin did not reopen a social reward learning critical period.

### Non-replicable persistent effects can be explained by a small or nonexistent effect size

The surprising lack of replicable persistent effects observed in this study could arise because: (1) the persistent effects of psilocybin are small and thus not able to be readily replicated, (2) methodological differences between labs preclude replicability, and/or (3) the positive results are spurious. To assess the role of small effect size in the replicability of our experiments, we first estimated the effect size of psilocybin across all experiments with linear mixed effects modeling (**Methods**). Consistent with our pooled datasets, psilocybin had a substantial acute effect on the head-twitch response and modest but significant acute effects on behaviors in the open field test, elevated plus maze, novel object exploration, and fear conditioning. Persistent effects were small across all experiments (**Extended Data Fig. 5a, b**). To assess the influence of psilocybin’s effect size on replicability, we estimated the probability of observing a significant main effect of psilocybin with simulation experiments (**Methods**). Persistent effects with small effect sizes had low replicability probabilities, which paled in comparison to the probabilities for the more robust, acute effects observed (**Extended Data Fig. 5c-e**).

Next, to explore whether mouse-, environmental-, and experimental-specific conditions systematically altered mouse behaviors, which could limit replicability, we added these conditions as fixed effects to our linear mixed effects models. In agreement with the well-established influence of environmental conditions on mouse phenotyping^60,61^, we found that methodological variables significantly influenced mouse behavior in several experiments (**Extended Data Fig. 6a**). For experiments with nonreplicable persistent effects (namely elevated plus maze, novel object exploration, forced swim test, and fear extinction), several methodological variables significantly influenced behavior, including handling, injection acclimation, apparatus dimensions, and mouse weight. However, these variables did not significantly influence psilocybin’s effect size, with the exception of handling on psilocybin’s effect in novel object exploration (**Extended Data Fig. 6b**), suggesting that overall, nonreplicable persistent effects were unlikely to result from the influence of methodological differences.

A final possibility which could explain the absence of replicable persistent effects is that isolated positive findings were spurious, false positives. When observing α=0.05, 5% of independent experiments are expected to be false positives, and indeed in our study of close to 80 total independent behavioral experiments, there were 4 occasions of singular, non-replicable positive effects. Together, these analyses suggest that nonreplicable persistent effects observed in this study are likely a result of psilocybin having a small or nonexistent persistent effect at the dose tested, rather than methodological differences.

## Discussion

We have presented results from a multi-institutional collaborative effort designed to elucidate the replicable acute and persistent effects of psilocybin on a range of mouse behaviors. Despite large quantitative variance in many of our behavioral measurements due to both lab-to-lab variability and individual subject variability, overall, many effects of psilocybin were replicable between labs (**Extended Data Fig. 7**). Specifically, we found that psilocybin acutely increased anxiety-related behaviors and decreased fear expression. Surprisingly, however, we found no evidence for replicable persistent effects of psilocybin 1-2 days after drug administration on anxiety-, depression-, and fear extinction-related behaviors. Furthermore, psilocybin did not reopen a critical period for social reward learning in adult subjects. These findings call into question the widespread notion that psilocybin promotes profound, long-lasting behavioral changes in commonly used behavioral assays. They also emphasize the advantages of this collaborative research effort and that, as many scientists appreciate but rarely explicitly state, results from any individual laboratory must be interpreted with caution.

The reliable acute anxiogenic effects of psilocybin in the OFT and EPM reported here are consistent with previous work on the effects of psilocybin in the open field using higher doses (3-5 mg/kg)^22,35,40^. They also are consistent with human subjects’ reports of psilocybin generating acute anxiety^3,4,62^. However, these results are inconsistent with those for the psychedelic 2,5-dimethoxy-4-iodoamphetamine (DOI), which has been reported to decrease avoidance behaviors in mice that were assessed using the elevated plus maze^63–65^ or marble burying test^63,66^. Additional work will be necessary to determine whether the inconsistent reported effects of psychedelics on anxiety-related behaviors are due to differences in the drugs being studied, methodological differences in the assays being performed, and/or generation of unreliable or spurious results.

A second acute effect of psilocybin that was remarkably replicable across labs was its effect on reducing fear expression. Every single lab observed this effect, and to our knowledge, every study in the literature has reported similar results^37,39,45^. Considering the replicability of this effect, future mechanistic studies of this effect are warranted, and if translatable to humans, this finding suggests the potential utility of psilocybin for suppressing fear responses during post-traumatic stress disorder exposure therapy.

It was surprising that replicable persistent effects of psilocybin on anxiety, depression, and fear extinction-related behaviors were not detected. It should be noted, however, that in several of these assays, one lab found statistically significant effects. We propose three possible interpretations for these isolated observations: (1) psychedelics may have some long-lasting effects on these behaviors, but that the true effect size is much smaller than previously anticipated and thus not able to be consistently replicated (**Extended Data Fig. 5**), (2) the results are spurious and reflect chance variation, or (3) the results are extremely dependent on idiosyncratic testing conditions. We do not think that differences in results between labs were due to differences in our experimental methods because overall, psilocybin’s effects varied minimally between labs (**Extended Data Fig. 7b**), and methodological variables did not significantly influence psilocybin’s effect size when non-replicable effects were observed – apart from handling on novel object exploration (**Extended Data Fig. 6**).

It is possible that experiments conducted using different behavioral assays or doses might yield different effects. For example, psilocybin (1.5 or 3 mg/kg IP) was reported to have anxiolytic effects in the novelty-suppressed feeding test^22,24^ but not in the open field test when assayed four hours after drug administration^22^. Furthermore, psilocybin’s effect of facilitating fear extinction was reported to have an unusual dose-dependence with maximal effects at 1 mg/kg and null effects at 2 mg/kg^39^, albeit another study reported significant effects using 2.5 mg/kg^37^.

The relevance of results using mice to the effects of psychedelics in human subjects remains to be determined. Although we observed that psilocybin’s persistent behavioral effects in standard laboratory behavioral paradigms are minimal in mice, it is possible that factors unique to the human psychedelic experience cannot be mimicked in mice and may be critical for the potential enduring therapeutic effects of psilocybin. Another possibility is that psilocybin has effects that specifically manifest in pathological states but are not observed in standard laboratory mice, which have not been subjected to interventions such as stress. Nevertheless, it seems very likely that there will continue to be an increase in efforts devoted to elucidating the mechanisms of psychedelics in experimentally tractable species such as mice^47,55,63,64,67,68^ and that the results from these studies will be used to interpret the therapeutic actions of psychedelics in human subjects. Given the typical trajectory of scientific discovery from individual investigators, the replicability of new findings will only slowly be realized. Furthermore, basing conclusions and future studies on unreliable findings will impede, not accelerate, scientific understanding and progress. Thus, we believe that the advantages and importance of collaborative research efforts of the type described here are self-evident. We hope that our efforts to define the replicable effects of psilocybin on a variety of mouse behaviors using a collaborative approach inspires others to adopt similar approaches. In this way, we believe we can make the most rapid and important advances to psychedelic science.

## METHODS

### Mice

Male and female C57BL/6J mice (Jackson Laboratory, stock number: 000664) were used for all experiments. Records of experimental mouse ages, weights, and housing numbers are detailed in **Supplementary Table 1**. Mice were maintained on the following light:dark cycles in each lab: Stanford – 12:12 (lights on 7am-7pm); Berkeley 1 – 14:10 (lights on 6am-8pm); Berkeley 2 – 12:12 (lights on 9pm-9am or 11pm-11am); UCSF 1 – 12:12 (lights on 6am-6pm); UCSF 2 – 12:12 (lights on 6am-6pm). Light:dark cycles were shifted to follow daylight savings in Stanford, Berkeley 2, and UCSF 1. Experimental procedures complied with animal care standards set forth by the National Institutes of Health and were approved by each institution’s Administrative Panel on Laboratory Animal Care.

### Psilocybin

Psilocybin powder (obtained from the National Institute of Drug Abuse) was diluted in 0.9% sterile saline to a concentration of 0.2 mg/mL. Psilocybin solution was then administered intraperitoneally (IP) at a volume of 10 mL/kg to achieve a dose of 2 mg/kg for all experiments.

### Liquid chromatography-high resolution mass-spectrometry (LC-HRMS)

#### Analyte standards

Psilocybin certified reference material was purchased from Cerilliant^Ⓡ^ P-097 1mL; 1.0 mg/mL in acetonitrile:water (1:1); lot#FE07232102. For use as an internal standard (IS), 5-Hydroxy-L-Tryptophan (5-HTP) was purchased from TCI America™ (≥98.0% purity determined by HPLC).

#### Calibration curve preparation

A psilocybin standard reference solution was serially diluted with saline to obtain a series of standard working solutions, and a 10 mM stock of 5-HTP in DMSO was diluted to give a 500 µg/mL IS working solution. The calibration curve was prepared by spiking 20 µL of each standard working solution into 200 µL water followed by addition of 20 µL IS. A calibration curve was prepared fresh with each set of samples and comprised a range of 0.5–50 µg/mL psilocybin in 200 µL aliquots.

#### Psilocybin Stability Samples Preparation

A sample of psilocybin used in the behavioral assays reported herein (0.2 mg/mL) was prepared in normal saline (0.9% NaCl, pH = 7.4, rt), aliquoted (200 uL) into triplicate 1.5 mL plastic tubes, and subjected to a variety of conditions: 1) freeze-thaw cycles (1x or 3x), 2) incubation at 4°C (3, 7, and 14 days), 3) incubation at 21°C under ambient 700 lux light (24 hours), 4) incubation at 21°C in the dark (8 or 24 hours), 5) incubation at pH = 10.05 and 21°C in the dark (100 mM sodium bicarbonate, 24 hours), or 6) incubation at pH = 2.05 and 21°C in the dark (0.1% formic acid, 24 hours). Samples for each condition were frozen at -80°C prior to analysis, whereupon they were thawed, vortexed, and diluted 20-fold in water followed by addition of 20 µL IS (500 µg/mL 5-HTP). Samples were then vortexed and centrifuged for 5 min at 10,000 rpm at 4°C, and 5 µl of supernatant was subjected to liquid-chromatography high resolution mass spectrometry (LC-HRMS).

#### LC-HRMS

A unified LC-UV-HRMS method was used to quantify psilocybin concentrations across sample conditions with a Waters Acquity H Class UPLC Plus Bio Quaternary system equipped with Orbitrap Exploris 240 (Thermo Fisher Scientific) HRMS. Liquid Chromatography was performed on an Atlantis® T3 (3 µm, 2.1 mm x 100 mm) column held at 40 °C using a linear gradient of mobile phases: A) 20 mM ammonium formate + 0.1% formic acid in water and B) 20 mM ammonium formate + 0.1% formic acid in acetonitrile/methanol/water 45:45:10. Elution was performed using an initial hold at 5% B for 3 minutes, followed by a gradient of 5–95% in 2 minutes, then held at 95% for 2 minutes; total run time was 10 minutes at a flow rate of 250 µL/min. Dual wavelength UV detection was performed at 254/269 nm and yielded retention times (RT) for psilocybin and 5-HTP of 2.79 min and 2.73 min, respectively.

Acquisition by HRMS was performed with heated electrospray ionization (HESI) in positive mode with full scan (m/z 100–600) at 120,000 orbitrap resolution. Data dependent (ddMS2) scanning in dynamic exclusion mode (isolation window 2m/z; normalized HCD collision energy=35v) was used to complement identification of parent compound and the internal standard. Product ion scans were collected to enhance method specificity, sensitivity and minimize background noise during detection. Precursor ions m/z were set up as follows: 285.0999 (psilocybin) and 221.0921 (5-HTP).

#### Quantification

Quantitative analysis was done with Thermo Xcalibur Quan Browser software (Thermo Fisher Scientific) using an internal standard approach. The mass transitions used for quantitation were as follows: psilocybin: m/z 285.0999 → m/z 205.1334 (quantifier); m/z 285.0999 → m/z 240.0419 (qualifier); and 5-HTP (IS): m/z 221.0921 → m/z 204.0653 (quantifier). Calibration curves were linear (R>0.98) over the concentration range using a weighting factor of 1/X^2^ where X is the concentration of psilocybin. The back-calculated standard concentrations were ±15% from nominal values, and ±20% at the lower limit of quantitation (LLOQ).

### Behavioral Assays

To promote replicability between labs, many factors that typically contribute to mouse behavioral variability were standardized. Namely, all labs performed experiments using male and female, adult, group-housed C57BL/6J mice. Prior to experiments, all mice were acclimated to the animal facility for at least one week, and mice were acclimated to the experimental room or a holding room for at least one hour. Prior to acute and persistent experiments, all labs handled (scruffed) and habituated mice to receiving injections (0.2 mL of 0.9% sterile saline IP or mock injection). For experiments testing the acute effects of psilocybin in the open field test and head twitch response assay, mice were tested immediately after injection. For all other experiments testing the acute effects of psilocybin, mice were injected with psilocybin (2 mg/kg) or saline and placed into clean, individual holding cages for 5-15 minutes before testing. For experiments testing the persistent effects of psilocybin, mice were injected with psilocybin (2 mg/kg) or saline and returned directly to their home cages for 24 hours before behavioral testing. For all experiments, behavioral equipment was cleaned with 70% EtOH before experiments, after experiments, and between individual animals. Experimenters at Stanford, Berkeley 1, Berkeley 2, and UCSF 2 were always blinded to the treatment group. For a more comprehensive report of experimental conditions for each mouse that could influence behavioral results (i.e., sexes, ages, housing numbers, dimensions of materials and apparatuses, handling counts, acclimation times, experiment times of day, number of mice tested concurrently, prior experimentation, and treatment details), see **Supplementary Table 1**.

### Head twitch response (HTR)

The head twitch response is a rapid, side-to-side, rotational head movement that is considered a proxy of hallucinogenic activity in mice^32,33^. To quantify psilocybin-induced head twitch responses, mice were injected with psilocybin or saline IP and monitored for 20 minutes. Head twitches were visually quantified by a blinded, trained observer.

### Open field test (OFT)

To assess psilocybin’s effects on locomotion and anxiety-like behaviors, mice were placed into an open field box and allowed to behave freely for 30 minutes (acute experiments) or 15-30 minutes (persistent experiments). Experiments were video recorded, and the average velocity (cm/s) and percent time spent in the center were automatically quantified using Ethovision, Biobserve, or DeepLabCut & custom MATLAB scripts. The center zone size varied from 39-57% of the total area of the open field.

### Elevated plus maze (EPM)

To assess psilocybin’s effects on anxiety/avoidance-like behaviors, mice were placed into the elevated plus maze, facing the open arm opposite the experimenter, and allowed to freely behave for 10 minutes for both acute and persistent experiments. Experiments were video recorded, and the percentage of time the animal’s center body was in the open arms was automatically computed using Ethovision, Biobserve, or DeepLabCut & custom MATLAB scripts.

### Social interaction test (SIT)

To assess psilocybin’s effects on social preference, mice underwent the social interaction test. In this test, two wire cups (to allow contact between mice) were placed into the left and right chambers of a three-chambered apparatus. Next, an ovariectomized female mouse or same-sex juvenile mouse was positioned inside one of the cups. Afterwards, the experimental mouse was placed into the center chamber of the three-chambered apparatus and allowed to explore freely for 10 minutes for both acute and persistent experiments. Experiments were video recorded, and the time exploring both the empty and social cups was quantified using automated methods (Ethovision, Biobserve, or DeepLabCut & custom MATLAB scripts). For all automated methods, an animal was considered exploring if its nose was within a zone defined around the cups. The social interaction index was defined as (time spent exploring the social cup – time spent exploring the empty cup) / (total time exploring both cups). Animals placed inside the cups were habituated to the cups prior to experiments, and experimental mice were habituated to the 3-chambered apparatus (with cups but no animals) prior to the experiment.

### Novel object exploration (NOE)

To assess psilocybin’s effects on exploration of a novel object, two identical objects were placed into an open field box. Then, mice were placed into the open field and allowed to explore freely for 10 minutes. Experiments were video recorded, and the total time spent exploring the objects was quantified using DeepLabCut & custom MATLAB scripts. The animal was considered exploring if its nose was within an object’s zone.

### Tail suspension test (TST)

To assess psilocybin’s persistent effects on passive coping behaviors in naïve mice, mice underwent the tail suspension test as previously described^57^. Briefly, mice were hung by their tails, at least 25 cm from the ground, for 6 minutes. Experiments were video recorded, and the time the animal spent immobile was quantified with Ethovision.

### Repeated tail suspension test / repeated forced swim test

To assess psilocybin’s persistent effects on passive coping behaviors in stressed mice, we used a protocol similar to one that has yielded antidepressant-like effects of psychedelics^28^. Specifically, for the repeated tail suspension test, mice were first handled for one to three consecutive days (1 minute per mouse) prior to experimentation (exception: UCSF 2). On day 1 of the experiment, mice underwent a 6-minute TST pre-test, following the protocol described above, to induce a depression-like phenotype. On day 2, mice were administered saline or psilocybin IP. On day 3, mice underwent a 6-minute TST post-test to assess whether psilocybin could reduce stress-induced increases in passive coping. All experiments were video recorded, and immobility was quantified using Ethovision.

The repeated forced swim test was conducted exactly as previously described^28^. Briefly, mice were first handled for one to three consecutive days (1 minute per mouse) prior to experimentation (exception: UCSF 1). On day 1 of the experiment, mice underwent a 6-minute FST pre-test, in which mice were placed in a clear Plexiglas cylinder filled with 24 ± 1°C water to induce a depression-like phenotype. On day 2, mice were administered saline or psilocybin IP. On day 3, mice underwent a 6-minute FST post-test to assess whether psilocybin could reduce stress-induced increases in passive coping. All experiments were video recorded, and immobility scores were determined via manual scoring by a trained, blinded observer.

### Cued fear conditioning

To assess psilocybin’s persistent effects on fear extinction learning, we performed fear conditioning experiments. On day 1 of the experiment (encoding session), mice were placed into context 1, which included a lemon scent, metal bar flooring and rectangular walls. The encoding session consisted of a 300 second habituation period, followed by five tone-shock pairings (30 seconds, 2.9 or 5 kHz, 65-75 dB pip tone, concluding with a 2 second, 0.75 mA footshock). The intertrial interval ranged from 55-65 seconds. On day 2 of the experiment (retrieval session), mice were placed into context 2, which included an anise scent, hard plastic flooring, and round walls. The retrieval session consisted of a 180 second habituation period, followed by 2 tones (same tones as during encoding), separated by a 50 second intertrial interval. On day 3 and 4 of the experiment (extinction sessions), mice were again placed into context 2. The extinction sessions consisted of a 180 second habituation period, followed by 10 tones (same tones as during encoding), with intertrial intervals ranging from 55 to 65 seconds. At the conclusion of each session, mice were removed from fear boxes ∼60 seconds after the final tone and returned to their home cage or holding cage. UCSF labs used holding cages to transfer mice between the holding room and experimental room. For injection before retrieval experiments, mice were injected 5-10 minutes before the retrieval session and placed into individual holding cages. For injection after retrieval experiments, mice were injected 5-10 minutes after the retrieval session and returned to their home cages directly after. All experiments were video recorded and analyzed by a blinded, trained observer or through automated methods (FreezeFrame or custom MATLAB scripts).

### Social conditioned place preference

To assess whether psilocybin could reopen the social reward learning critical period, mice underwent a social conditioned place preference task. In this task, mice were first single-housed for three days prior to any experimentation to prime a desire for social interaction^47^. On day 1 of the social conditioned place preference experiment, mice underwent a 30- or 60-minute pre-test, during which mice were placed into skinner boxes or custom-made boxes that had two distinct contexts on the left and right sides. Textured / smooth flooring and striped / polka-dot walls were used to create distinct contexts. Then, on days 2-4, mice experienced 1-hour periods of social conditioning, in which they were re-united with 1-4 cage mates in one context, and 1-hour periods of isolation conditioning, in which they were alone in the other context. Finally, on day 5, mice underwent a 30- or 60-minute post-test, in which they had free access to explore both contexts in the skinner boxes. To quantify social reward learning, we identified the mouse’s body center using automated methods (Biobserve, Ethovision, or DeepLabCut) and compared time spent in the social context during the pre-test versus post-test. The social preference score was calculated as the time spent in the social context during the post-test divided by the time spent in the social context during the pre-test. Mice that exhibited strong baseline preferences for one chamber, defined as spending more than 75 percent of time in one chamber during the pre-test, were excluded.

Using this task, each lab first tested whether juvenile mice (P34-41) experienced social reward learning, defined as having a social preference score that was significantly different from 1.0 (a value indicative of no social reward learning). In labs that established social reward learning in juvenile mice, adult mice (P88-106) were then tested to determine if, as previously reported^47^ social reward learning was absent at this age. To test whether psilocybin could ‘reopen’ the social reward learning critical period in adult mice, mice were pre-treated with saline or psilocybin four days prior to the start of experiments (while mice were still group housed, 1 day prior to single-housing).

### Simulations

To determine the distribution of p-values expected if 10,000 labs were to repeat our experiments, we first z-scored each lab’s data to each lab’s saline data, for males and females separately to determine the normalized size of the effect in each experiment. Next, we resampled a kernel density estimation of our z-scored data 10,000 times and used 2-way ANOVAs (factors: drug and sex) or 3-way ANOVAs (for fear conditioning; factors drug, sex, and cue number) to analyze each sample. The resulting distribution of p-values for the main drug effect was then plotted. The replicability probability was calculated for experiments in which at least one lab showed a positive result and was defined as the percentage of experiments expected to yield a p-value less than 0.05.

### Statistical Analyses

#### Two-way ANOVAs

To determine whether psilocybin had a significant effect on mouse behavior, and whether there were any sex-specific effects, we performed two-way ANOVAs using Prism 10 (Graphpad), with drug and sex as factors. Significance for main effects of drug and/or sex are visualized with asterisk(s) next to either the drug legend or next to sex conditions, respectively. When there was a significant interaction between drug treatment and sex, Sidak’s multiple comparisons were used to test for any sex-specific effects of the drug, and significance from post-hoc comparisons is denoted on the main body of the graphs. Statistical significance was *p<0.05, **p<0.01, ***p<0.001. Data are presented as mean ± SEM.

#### Repeated measures three-way ANOVAs

Repeated measures three-way ANOVAs were performed when there was an additional third variable of interest (i.e., timepoint, cue number, object type, cup type). Factors included drug, sex, the third variable of interest, and all interactions. Statistical significance was *p<0.05, **p<0.01, ***p<0.001. Data are presented as mean ± SEM.

#### Linear mixed effects modeling

The percentage of variance explained by psilocybin treatment (**Extended Data Fig. 5a**) was determined through a linear mixed-effects model (R version 4.4.1, package lme4^69^) of the raw pooled data, with a fixed effect of psilocybin treatment and random effects of lab and cage. Models were fit separately for each experiment, and the following outcome variables were used: number of head twitch responses in 20 minutes (HTR), % time spent in the center of the open field during acute (aOFT - center) and persistent (pOFT - center) open field test experiments, average velocity (cm/s) during the acute (aOFT – velocity) and persistent (pOFT – velocity) open field test experiments, % time spent in the open arms of the elevated plus maze during acute (aEPM) and persistent (pEPM) experiments, average exploration time (s) of novel objects during acute (aNOE) and persistent (pNOE) experiments, social preference index from acute (aSIT) and persistent (pSIT) social interaction test experiments, day 3 immobility times (s) from the repeated forced swim test (rFST) and repeated tail suspension test (rTST), immobility times (s) from the persistent tail suspension test (pTST) in stress-naïve mice, average % time freezing to cues during retrieval for fear conditioning injection before retrieval experiments (FC – pre – retr), average % time freezing to cues during the first extinction session for fear conditioning injection before retrieval (FC – pre – ext1) and injection after retrieval (FC – post – ext1) experiments, average % time freezing to cues during the second extinction session for fear conditioning injection before retrieval (FC – pre – ext2) and injection after retrieval (FC – post – ext2) experiments, and social preference scores for adult saline versus psilocybin social conditioned place preference (sCPP) experiments. The abbreviated names in parentheses are what is presented in the extended data figures.

To determine psilocybin’s effect size in z-score (**Extended Data Fig. 5b**), we first z-score normalized all data points in each lab to each lab’s saline groups for each sex separately before fitting linear mixed effects models. Outcome variables used for each model are as described above. Model statistics were evaluated using R packages MuMIn^70^ and lmerTest^71^.

To determine fixed effects due to treatment or methodology and random effects due to lab or cage, we used additional linear mixed-effects models. Fixed effects, with abbreviated names in brackets, included: i. treatment given: saline or psilocybin [treatment psi], ii. mouse sex [sex male], iii. mouse age in weeks [age], iv. mouse weight in grams [weight], v. number of handling acclimation sessions [handling acc.], vi. number of injection acclimation sessions [injection acc.], vii. number of weeks of acclimation to the lab [lab acc.], viii. number of cage mates [cagemates], ix. same treatments within a cage [yes=1, no=0] [same treatment], x. timepoint of testing after treatment [timepoint], xi. number of simultaneous other mice tested [other mice], xii. number of prior experiments [prior expts], xiii. apparatus dimensions [dimension 1; dimension 2], and xiv. time of experiment relative to light-cycle [time of day]. Features that had the same value for all mice tested across the five labs were excluded from the model. The fixed effect size of each feature in each experiment was estimated with 95% confidence intervals. Outcome variables used for each model are as described above. For the methodological variables that had a significant influence on mouse behavior in non-replicable persistent experiments, simple linear regression was performed to quantify the effect of that variable on psilocybin’s effect size, where x = selected methodological variables, and y = z-scored psilocybin data for the selected experiment. Slope values from linear regression were estimated with 95% confidence intervals.

To assess how mouse behavioral outcomes depended on the laboratory in which they were conducted, we evaluated the degree of clustering by comparing the intraclass correlation (ICC) across all behavioral paradigms. ICC was calculated for both the raw data to identify the inter-lab variability in the raw measurements (**Extended Data Fig. 7a**), as well as for the z-score normalized data to identify the inter-lab variability in psilocybin’s effect (**Extended Data Fig. 7b**).

## DATA AVAILABILITY

Source data will be made available with this paper.

## CODE AVAILABILITY

All custom code used for analysis in this manuscript is publicly available on Github: https://github.com/odilialu/Psy-BAANC.

## ACKNOWLEDGEMENTS

This work was philanthropically funded by a donor who wishes to remain anonymous. Portions of this work were also supported by P50 DA042012 (BDH and RCM). OL was supported by an Alper Fellowship in the Neurosciences. CL was supported by a UCSF NIH IRACDA Fellowship (K12GM081266). MBP was supported by NIH grant K99 DA056573. HRMS experiments were supported by the Vincent Coates Foundation Mass Spectrometry Laboratory, Stanford University Mass Spectrometry (RRID:SCR_017801). HRMS data were collected using a Thermo Exploris 240 LC/MS system (RRID:SCR_022216) purchased with funding support from the Community of Shared Research Platforms at Stanford University.

## AUTHOR CONTRIBUTIONS

Behavioral experiments: Head twitch response quantifications were completed by KR, RL, MZ, KW, AK, AG. Open field tests were conducted by ODL, KR, KW, NG, CL, RL, AL. Elevated plus maze experiments were conducted by ODL, KR, KW, NG, AG. Novel object exploration experiments were conducted by ODL, KR, KW, NG, LS, AL. Social interaction tests were conducted by ODL, KR, KW, NG, CL, AL. Tail suspension tests were conducted by MZ, KW, NG, SV, LS, AL, CL. Forced swim tests were conducted by RL, MZ, KW, AK, LS, AL, SV. Fear conditioning experiments were conducted by RL, MZ, KW, NG, SV, CL. Analyses were completed by ODL, KW, NG, KR, SV, CL, AK, AG, LS, AL. Supervision by ODL, ABC, LPC, MBP, AK, CL. Consortium coordination by ODL. Simulations by ODL. Mixed modeling by KW. Conceptualization by VS, MAK, AG, SL, BH, RM. Manuscript written by ODL, SL, RM. Figures composed by ODL. All authors contributed to editing the final version of this manuscript.

## ADDITIONAL INFORMATION

Correspondence and requests for materials should be addressed to Robert C. Malenka.

## FIGURE LEGENDS

**Extended Data Figure 1.**
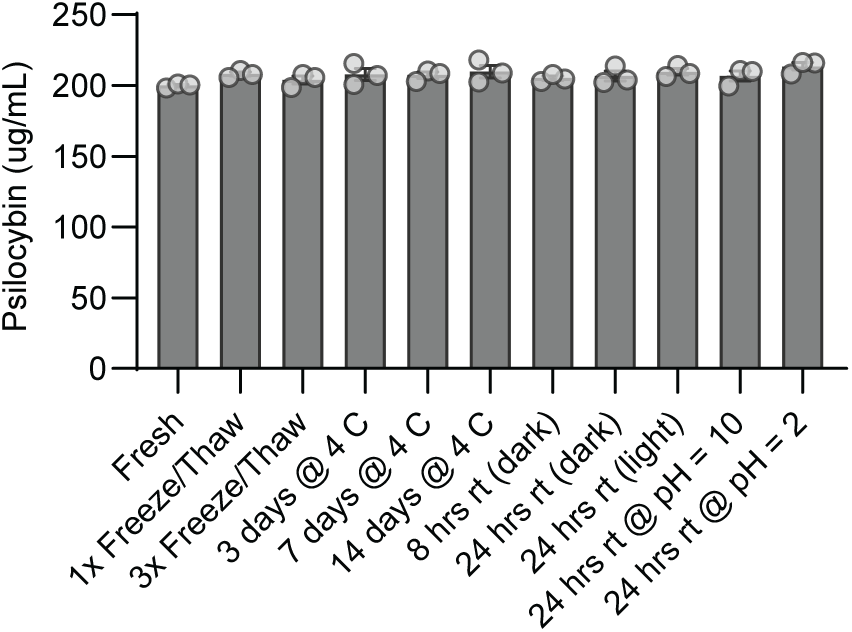
2 mg/kg psilocybin in saline preparation is stable across multiple conditions. Bar graph showing high resolution mass spectrometry data indicating that psilocybin dissolved in saline is stable across multiple freeze-thaw cycles, different temperatures (rt = room temperature), lighting conditions, and acidity levels (n=3 per condition). Data shown as mean±SEM. Data analyzed with ordinary one-way ANOVA (F=1.431, p=0.2313). Detailed statistics for all analyses are provided in **Supplementary Table 2**.

**Extended Data Figure 2.**
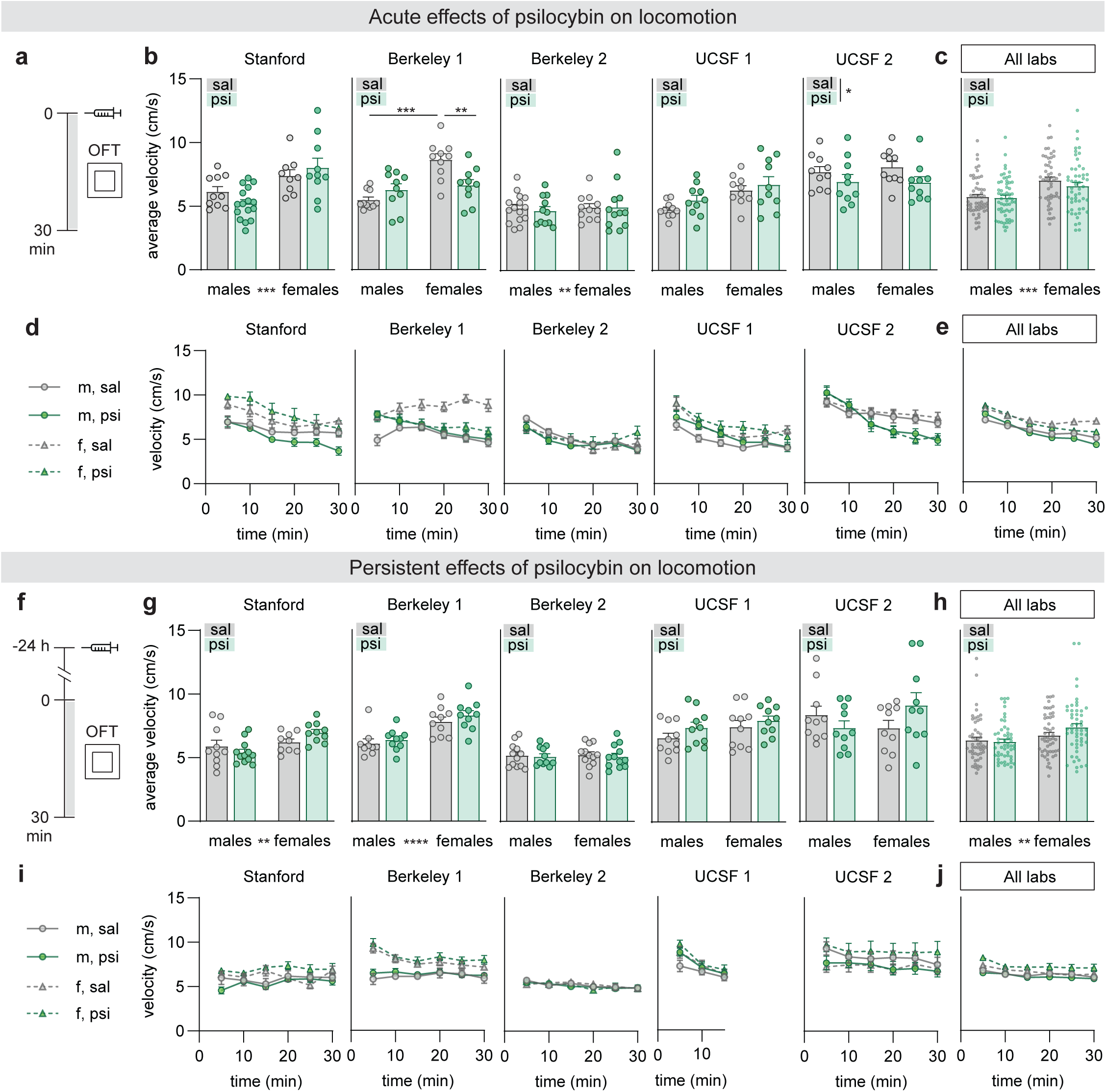
Psilocybin has no reproducible acute or persistent effects on locomotion. **(a)** Experimental timeline to test the acute effects of psilocybin on locomotion in the open field test (OFT). **(b)** Average mouse velocity over the 30 minute OFT for each individual lab. **(c)** Data from panel b pooled together. **(d)** Time course analysis of velocity (5-min time bins). **(e)** Data from panel d pooled together. **(f-j)** As in panels a-e, but for experiments testing the persistent effects of psilocybin on locomotion. Data shown as mean+SEM. *p<0.05, **p<0.01, ***p<0.001. Detailed statistics are provided in **Supplementary Table 2**.

**Extended Data Figure 3.**
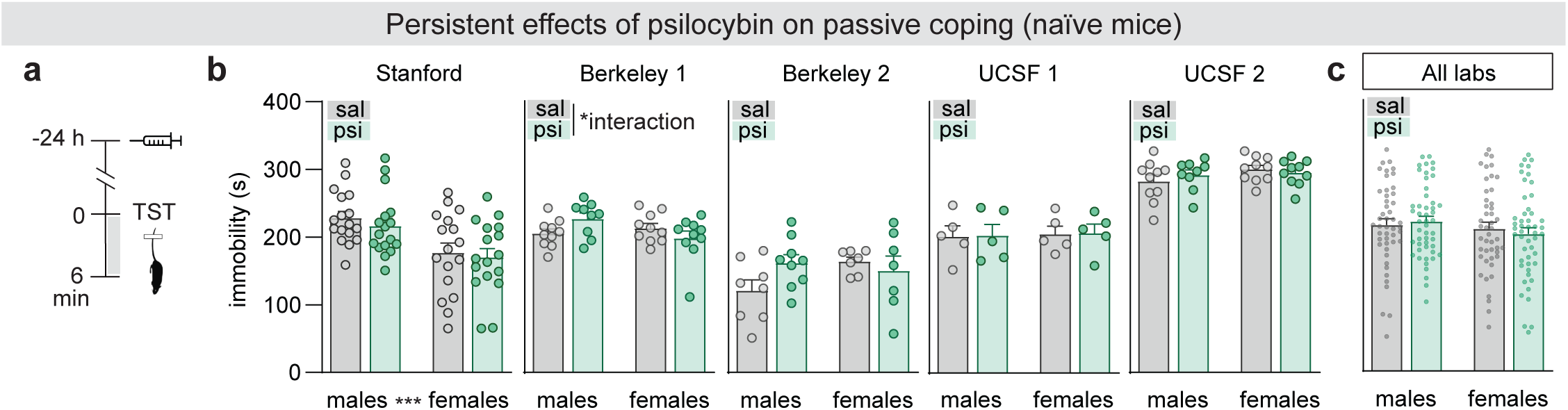
Psilocybin has no effects on depression-related behaviors in naive mice. **(a)** Experimental timeline to test the persistent effects of psilocybin on depression-related behaviors in the tail suspension test (TST). **(b)** Time mice spent immobile in the TST for each individual lab. **(c)** Data in panel b pooled together. Data shown as mean+SEM. *p<0.05, **p<0.01, ***p<0.001. Detailed statistics are provided in **Supplementary Table 2**.

**Extended Data Figure 4.**
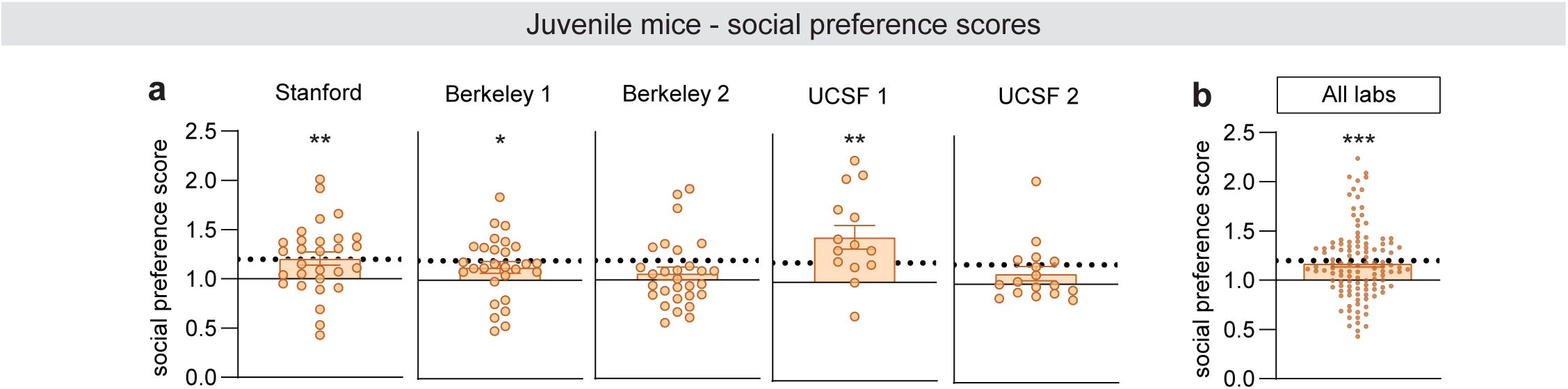
Social conditioned place preference scores for juvenile mice in each individual lab. **(a)** Bar graphs plotting social preference scores in the social conditioned place preference experiment for juvenile mice in each individual lab. **(b)** Data in panel a pooled together. Dotted line at y = 1.2 indicates the social preference score expected of juvenile mice based on the literature (Nardou et al., 2019, 2023). Stanford, Berkeley 1, and UCSF 1 juvenile mice exhibited a significant social conditioned place preference (one-sample t-test; hypothetical mean = 1.0). Data shown as mean±SEM. *p<0.05, **p<0.01, ***p<0.01. Detailed statistics are provided in **Supplementary Table 2**.

**Extended Data Figure 5.**
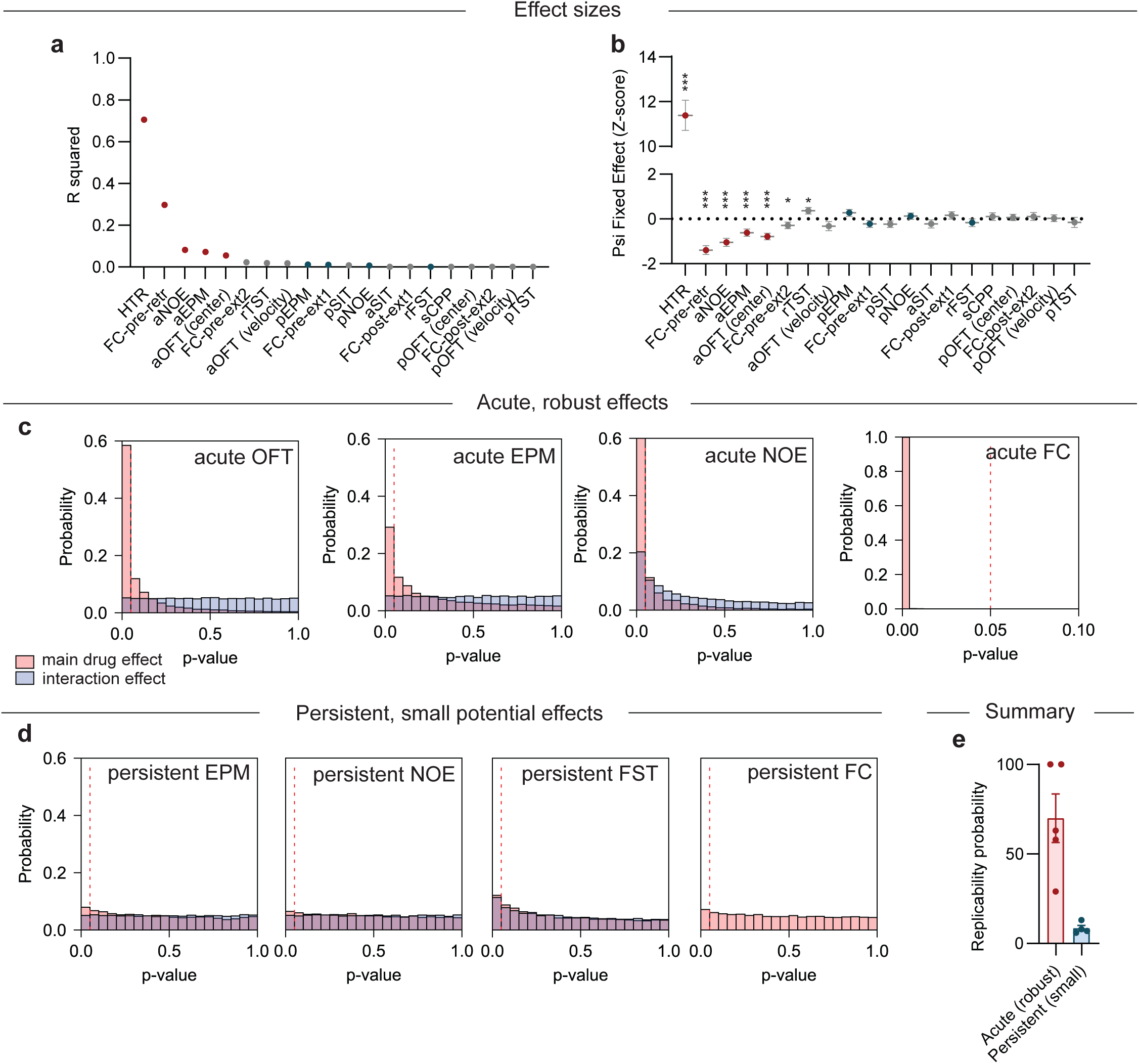
Robust effects are easily reproduced across labs, whereas small effects are challenging to replicate. **(a)** Proportion of variance in data explained by psilocybin treatment across all behaviors tested indicates effect size. Mixed effects models included psilocybin treatment as a fixed effect and lab and cage as random effects. Experiments where at least one lab showed a significant effect have colored points, with acute effects in red and persistent effects in blue. **(b)** Effect size of psilocybin across experiments, based on data Z-scored to the saline conditions for each experiment. Data presented as mean±SEM. *p<0.05, ***p<0.001. **(c)** Histogram of the distribution of p-values expected for main drug effect and interaction effect obtained from 10,000 simulated experiments based on acute OFT, acute EPM, acute NOE, and acute FC data, from left to right. **(d)** Histogram of the distribution of p-values expected for main drug effect and interaction effect obtained from 10,000 simulated experiments based on persistent EPM, persistent NOE, persistent FST, and persistent FC data, from left to right. **(e)** Bar graph comparing replicability probability from simulation experiments of acute, robust effects versus persistent, small effects. Only experiments where at least one lab showed an effect of psilocybin were included. Model statistics and details are available in **Supplementary Table 2** and **Methods.**

**Extended Data Figure 6.**
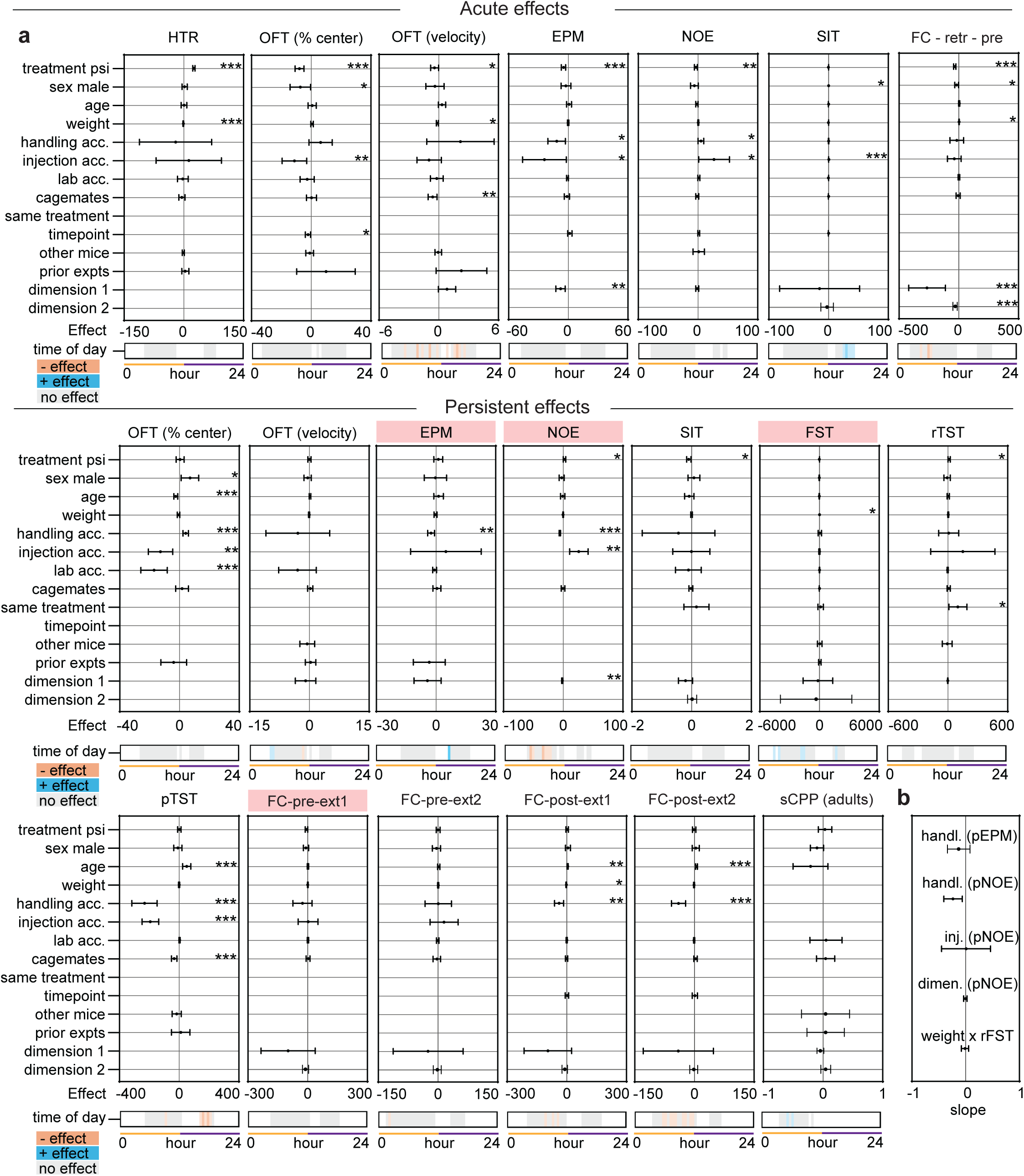
Mixed modeling analyses characterize the influence of methodological variables on behavior. **(a)** Mixed modeling analysis with methodological variables included as fixed effects. Experiments with a non-replicable persistent effect have a red-highlighted title. **(b)** Effect of the selected methodological variable on psilocybin’s effect in z-score. Data shown as mean±95% confidence intervals. *p<0.05, **p<0.01, ***p<0.001. Detailed descriptions of all fixed effects are available in the methods. Raw data used for modeling are available in **Supplementary Table 1**, and statistics are available in **Supplementary Table 2**.

**Extended Data Figure 7.**
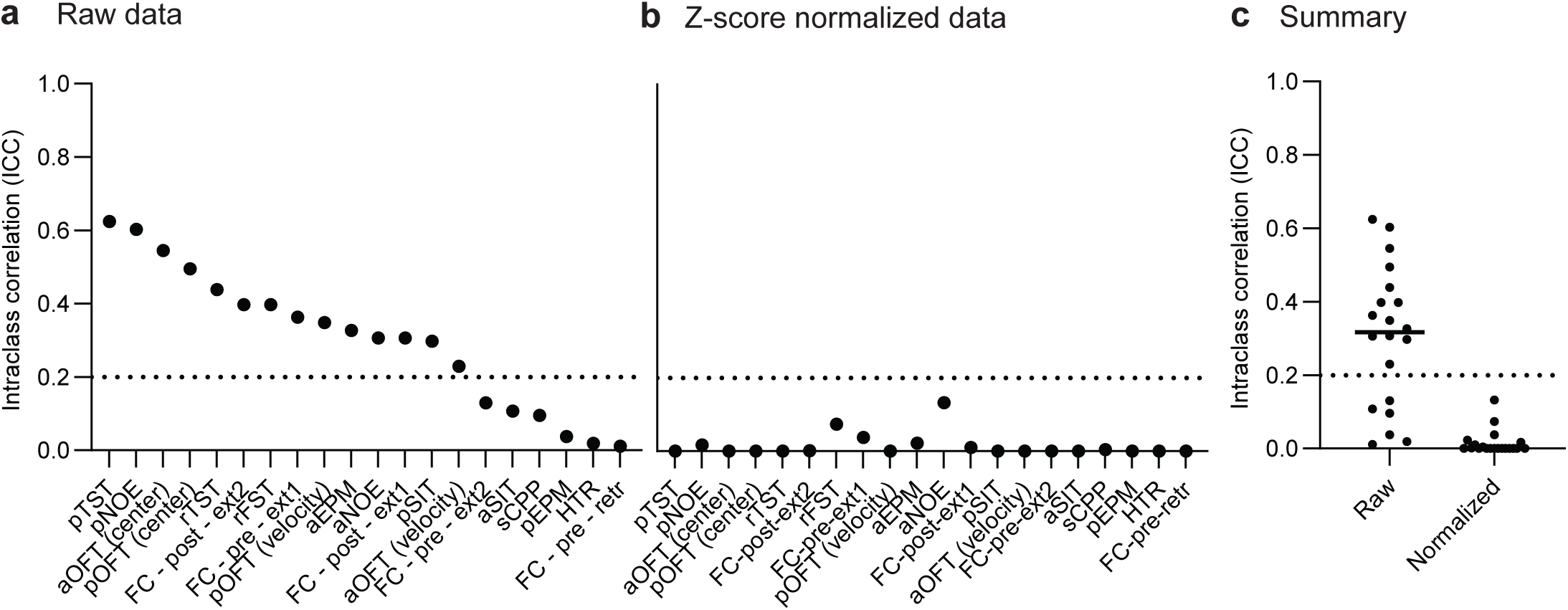
Despite large variance in raw data between labs, the effect of psilocybin was generally consistent across labs. **(a)** Intraclass correlation for each behavioral test shows large lab-to-lab variance. **(b)** Intraclass correlation after z-score normalizing data to saline data shows minimal lab-to-lab variance, indicating that effects of psilocybin varied minimally across labs. **(c)** Summary visualization of panels a and b. Dotted line at ICC = 0.2 indicates the value that most pragmatic cluster-randomized trials fall below. Model statistics are available in **Supplementary Table 2.**

## REFERENCES

1. Carhart-Harris, R. L. et al. Psilocybin with psychological support for treatment-resistant depression: an open-label feasibility study. Lancet Psychiatry 3, 619–627 (2016).

2. Raison, C. L. et al. Single-Dose Psilocybin Treatment for Major Depressive Disorder: A Randomized Clinical Trial. JAMA 330, 843 (2023).

3. Goodwin, G. M. et al. Single-Dose Psilocybin for a Treatment-Resistant Episode of Major Depression. N. Engl. J. Med. 387, 1637–1648 (2022).

4. Griffiths, R. R. et al. Psilocybin produces substantial and sustained decreases in depression and anxiety in patients with life-threatening cancer: A randomized double-blind trial. J. Psychopharmacol. (Oxf.) 30, 1181–1197 (2016).

5. Ross, S. et al. Rapid and sustained symptom reduction following psilocybin treatment for anxiety and depression in patients with life-threatening cancer: a randomized controlled trial. J. Psychopharmacol. (Oxf.) 30, 1165–1180 (2016).

6. Johnson, M. W., Garcia-Romeu, A., Cosimano, M. P. & Griffiths, R. R. Pilot study of the 5-HT 2A R agonist psilocybin in the treatment of tobacco addiction. J. Psychopharmacol. (Oxf.) 28, 983–992 (2014).

7. Bogenschutz, M. P. et al. Psilocybin-assisted treatment for alcohol dependence: A proof-of-concept study. J. Psychopharmacol. (Oxf.) 29, 289–299 (2015).

8. Von Rotz, R. et al. Single-dose psilocybin-assisted therapy in major depressive disorder: a placebo-controlled, double-blind, randomised clinical trial. eClinicalMedicine 56, 101809 (2023).

9. Mitchell, J. M. et al. MDMA-assisted therapy for severe PTSD: a randomized, double-blind, placebo-controlled phase 3 study. Nat. Med. 27, 1025–1033 (2021).

10. Mitchell, J. M. et al. MDMA-assisted therapy for moderate to severe PTSD: a randomized, placebo-controlled phase 3 trial. Nat. Med. 29, 2473–2480 (2023).

11. A Phase 2, Multi-center, Randomized, Double-Blind, Parallel-Group, Dose-Finding Study to Assess the Effect of Four Doses of MM-120 (LSD D-Tartrate) for the Treatment of Anxiety Symptoms. (2022).

12. Gasser, P. et al. Safety and Efficacy of Lysergic Acid Diethylamide-Assisted Psychotherapy for Anxiety Associated With Life-threatening Diseases. J. Nerv. Ment. Dis. 202, 513–520 (2014).

13. Cherian, K. N. et al. Magnesium–ibogaine therapy in veterans with traumatic brain injuries. Nat. Med. 30, 373–381 (2024).

14. Noller, G. E., Frampton, C. M. & Yazar-Klosinski, B. Ibogaine treatment outcomes for opioid dependence from a twelve-month follow-up observational study. Am. J. Drug Alcohol Abuse 44, 37–46 (2018).

15. Davis, A. K., Xin, Y., Sepeda, N. & Averill, L. A. Open-label study of consecutive ibogaine and 5-MeO-DMT assisted-therapy for trauma-exposed male Special Operations Forces Veterans: prospective data from a clinical program in Mexico. Am. J. Drug Alcohol Abuse 49, 587–596 (2023).

16. COMPASS Pathways. COMPASS Pathways receives FDA Breakthrough Therapy designation for psilocybin therapy for treatment-resistant depression. (2018).

17. Usona Institute. FDA grants Breakthrough Therapy Designation to Usona Institute’s psilocybin program for major depressive disorder. (2019).

18. MindMed. MindMed Receives FDA Breakthrough Therapy Designation and Announces Positive 12-Week Durability Data From Phase 2B Study of MM120 for Generalized Anxiety Disorder. (2024).

19. MAPS. FDA Grants Breakthrough Therapy Designation for MDMA-Assisted Therapy for PTSD, Agrees on Special Protocol Assessment for Phase 3 Trials. (2017).

20. Carhart-Harris, R. et al. Trial of Psilocybin versus Escitalopram for Depression. N. Engl. J. Med. 384, 1402–1411 (2021).

21. Hendrie, C. & Pickles, A. Psilocybin: panacea or placebo? Lancet Psychiatry 3, 805–806 (2016).

22. Jones, N. T. et al. Transient Elevation of Plasma Glucocorticoids Supports Psilocybin-Induced Anxiolysis in Mice. ACS Pharmacol. Transl. Sci. 6, 1221–1231 (2023).

23. Hesselgrave, N., Troppoli, T. A., Wulff, A. B., Cole, A. B. & Thompson, S. M. Harnessing psilocybin: antidepressant-like behavioral and synaptic actions of psilocybin are independent of 5-HT2R activation in mice. Proc. Natl. Acad. Sci. 118, e2022489118 (2021).

24. Takaba, R. et al. Ethopharmacological evaluation of antidepressant-like effect of serotonergic psychedelics in C57BL/6J male mice. Naunyn. Schmiedebergs Arch. Pharmacol. 397, 3019–3035 (2024).

25. Thakur, N. Investigating the Effects of Psilocybin on Models of Anxiety, Recognition Memory, and Depression-Like Behavior and the Role of the 5-HT2A Receptor in Mediating Psilocybin’s Impact on Behavioral Despair. doi:10.25772/02WQ-AJ35.

26. Jefsen, O. et al. Psilocybin lacks antidepressant-like effect in the Flinders Sensitive Line rat. Acta Neuropsychiatr. 31, 213–219 (2019).

27. Sekssaoui, M., Bockaert, J., Marin, P. & Bécamel, C. Antidepressant-like effects of psychedelics in a chronic despair mouse model: is the 5-HT2A receptor the unique player? Neuropsychopharmacology 49, 747–756 (2024).

28. Cameron, L. P. et al. 5-HT2ARs Mediate Therapeutic Behavioral Effects of Psychedelic Tryptamines. ACS Chem. Neurosci. 14, 351–358 (2023).

29. Saré, R. M., Lemons, A. & Smith, C. B. Behavior Testing in Rodents: Highlighting Potential Confounds Affecting Variability and Reproducibility. Brain Sci. 11, 522 (2021).

30. Laraway, S., Snycerski, S., Pradhan, S. & Huitema, B. E. An Overview of Scientific Reproducibility: Consideration of Relevant Issues for Behavior Science/Analysis. *Perspect*. Behav. Sci. 42, 33–57 (2019).

31. Hibicke, M., Landry, A. N., Kramer, H. M., Talman, Z. K. & Nichols, C. D. Psychedelics, but Not Ketamine, Produce Persistent Antidepressant-like Effects in a Rodent Experimental System for the Study of Depression. ACS Chem. Neurosci. 11, 864–871 (2020).

32. Corne, S. J. & Pickering, R. W. A possible correlation between drug-induced hallucinations in man and a behavioural response in mice. Psychopharmacologia 11, 65–78 (1967).

33. Halberstadt, A. L., Chatha, M., Klein, A. K., Wallach, J. & Brandt, S. D. Correlation between the potency of hallucinogens in the mouse head-twitch response assay and their behavioral and subjective effects in other species. Neuropharmacology 167, 107933 (2020).

34. Passie, T., Seifert, J., Schneider, U. & Emrich, H. M. The pharmacology of psilocybin. Addict. Biol. 7, 357–364 (2002).

35. Fadahunsi, N. et al. Acute and long-term effects of psilocybin on energy balance and feeding behavior in mice. Transl. Psychiatry 12, 330 (2022).

36. Shao, L.-X. et al. Psilocybin induces rapid and persistent growth of dendritic spines in frontal cortex in vivo. Neuron 109, 2535–2544.e4 (2021).

37. Du, Y. et al. Psilocybin facilitates fear extinction in mice by promoting hippocampal neuroplasticity. Chin. Med. J. (Engl.) 136, 2983–2992 (2023).

38. Rijsketic, D. R. et al. UNRAVELing the synergistic effects of psilocybin and environment on brain-wide immediate early gene expression in mice. Neuropsychopharmacology 48, 1798–1807 (2023).

39. Woodburn, S. C., Levitt, C. M., Koester, A. M. & Kwan, A. C. Psilocybin Facilitates Fear Extinction: Importance of Dose, Context, and Serotonin Receptors. ACS Chem. Neurosci. 15, 3034–3043 (2024).

40. Harari, R., Chatterjee, I., Getselter, D. & Elliott, E. Psilocybin induces acute anxiety and changes in amygdalar phosphopeptides independently from the 5-HT2A receptor. iScience 27, 109686 (2024).

41. Halberstadt, A. L. & Geyer, M. A. Effect of Hallucinogens on Unconditioned Behavior. in Behavioral Neurobiology of Psychedelic Drugs (eds. Halberstadt, A. L., Vollenweider, F. X. & Nichols, D. E.) vol. 36 159–199 (Springer Berlin Heidelberg, Berlin, Heidelberg, 2016).

42. Geyer, M. A. et al. A characteristic effect of hallucinogens on investigatory responding in rats. Psychopharmacology (Berl.) 65, 35–40 (1979).

43. De Gregorio, D., et al. Lysergic acid diethylamide (LSD) promotes social behavior through mTORC1 in the excitatory neurotransmission. Proc. Natl. Acad. Sci. 118, e2020705118 (2021).

44. Mollinedo-Gajate, I. et al. Psilocybin rescues sociability deficits in an animal model of autism. Preprint at 10.1101/2020.09.09.289348 (2020).

45. Rogers, S. A., Heller, E. A. & Corder, G. Psilocybin-enhanced fear extinction linked to bidirectional modulation of cortical ensembles. Preprint at 10.1101/2024.02.04.578811 (2024).

46. Catlow, B. J., Song, S., Paredes, D. A., Kirstein, C. L. & Sanchez-Ramos, J. Effects of psilocybin on hippocampal neurogenesis and extinction of trace fear conditioning. Exp. Brain Res. 228, 481–491 (2013).

47. Nardou, R. et al. Psychedelics reopen the social reward learning critical period. Nature 618, 790–798 (2023).

48. Nardou, R. et al. Oxytocin-dependent reopening of a social reward learning critical period with MDMA. Nature 569, 116–120 (2019).

49. Seibenhener, M. L. & Wooten, M. C. Use of the Open Field Maze to Measure Locomotor and Anxiety-like Behavior in Mice. J. Vis. Exp. 52434 (2015) doi:10.3791/52434.

50. Walf, A. A. & Frye, C. A. The use of the elevated plus maze as an assay of anxiety-related behavior in rodents. Nat. Protoc. 2, 322–328 (2007).

51. Maner, J. K. & Schmidt, N. B. The Role of Risk Avoidance in Anxiety. Behav. Ther. 37, 181–189 (2006).

52. Yang, M., Silverman, J. L. & Crawley, J. N. Automated Three-Chambered Social Approach Task for Mice. Curr. Protoc. Neurosci. 56, (2011).

53. Can, A. et al. The Mouse Forced Swim Test. J. Vis. Exp. 3638 (2011) doi:10.3791/3638.

54. Cameron, L. P. et al. A non-hallucinogenic psychedelic analogue with therapeutic potential. Nature 589, 474–479 (2021).

55. Vargas, M. V. et al. Psychedelics promote neuroplasticity through the activation of intracellular 5-HT2A receptors. Science 379, 700–706 (2023).

56. Dong, C. et al. Psychedelic-inspired drug discovery using an engineered biosensor. Cell 184, 2779–2792.e18 (2021).

57. Can, A. et al. The Tail Suspension Test. J. Vis. Exp. 3769 (2011) doi:10.3791/3769.

58. Bardo, M. T. & Bevins, R. A. Conditioned place preference: what does it add to our preclinical understanding of drug reward? Psychopharmacology (Berl.) 153, 31–43 (2000).

59. Tzschentke, T. M. REVIEW ON CPP: Measuring reward with the conditioned place preference (CPP) paradigm: update of the last decade. Addict. Biol. 12, 227–462 (2007).

60. Crabbe, J. C., Wahlsten, D. & Dudek, B. C. Genetics of Mouse Behavior: Interactions with Laboratory Environment. Science 284, 1670–1672 (1999).

61. Kafkafi, N. et al. Reproducibility and replicability of rodent phenotyping in preclinical studies. Neurosci. Biobehav. Rev. 87, 218–232 (2018).

62. Yerubandi, A. et al. Acute Adverse Effects of Therapeutic Doses of Psilocybin: A Systematic Review and Meta-Analysis. *JAMA Netw*. Open 7, e245960 (2024).

63. Muir, J. et al. Isolation of psychedelic-responsive neurons underlying anxiolytic behavioral states. Science 386, 802–810 (2024).

64. Tiwari, P. et al. Ventral hippocampal parvalbumin interneurons gate the acute anxiolytic action of the serotonergic psychedelic DOI. Neuron 112, 3697–3714.e6 (2024).

65. Nic Dhonnchadha, B. Á., Bourin, M. & Hascoët, M. Anxiolytic-like effects of 5-HT2 ligands on three mouse models of anxiety. Behav. Brain Res. 140, 203–214 (2003).

66. Odland, A. U., Jessen, L., Kristensen, J. L., Fitzpatrick, C. M. & Andreasen, J. T. The 5-hydroxytryptamine 2A receptor agonists DOI and 25CN-NBOH decrease marble burying and reverse 8-OH-DPAT-induced deficit in spontaneous alternation. Neuropharmacology 183, 107838 (2021).

67. Moliner, R. et al. Psychedelics promote plasticity by directly binding to BDNF receptor TrkB. Nat. Neurosci. 26, 1032–1041 (2023).

68. Duque, M. et al. Ketamine induces plasticity in a norepinephrine-astroglial circuit to promote behavioral perseverance. Neuron S0896627324008365 (2024) doi:10.1016/j.neuron.2024.11.011.

69. Bates, D., Mächler, M., Bolker, B. & Walker, S. Fitting Linear Mixed-Effects Models Using **lme4**. J. Stat. Softw. 67, (2015).

70. Kamil Bartoń. MuMIn: Multi-Model Inference. 1.47.5 10.32614/CRAN.package.MuMIn (2010).

71. Kuznetsova, A., Brockhoff, P. B. & Christensen, R. H. B. **lmerTest** Package: Tests in Linear Mixed Effects Models. J. Stat. Softw. 82, (2017).

